# Microbiota of the sulfur cycle in an extremely contaminated Technosol undergoing pedogenesis: A culture-dependent and metagenomic approach

**DOI:** 10.1101/2023.12.06.570440

**Authors:** K. Demin, T. Minkina, S. Sushkova, Y. Delegan, Y. Kocharovskaya, A. Gorovtsov

**Affiliations:** Southern Federal University, 344090 Rostov-on-Don, Russia; G.K. Skryabin Institute of Biochemistry and Physiology of Microorganisms, Pushchino, 142290 Moscow, Russia

**Keywords:** soil microbiome, sulfur cycle, sulfate-reducing bacteria, metagenome-assembled genomes, Technosol, pollution

## Abstract

Understanding the microbial communities involved in the global sulfur cycle is crucial for comprehending key biogeochemical processes on Earth. However, most studies tend to focus on marine ecosystems, while investigations into the terrestrial sulfur cycle are scarce. In this study, we employed culture-dependent techniques and metagenomics to characterize sulfur-cycling microbiota in extremely contaminated soils. We analyzed shotgun and amplicon sequencing data to assess taxonomical diversity, metagenome-assembled genomes (MAGs) for functional diversity, and also calculated the most probable numbers (MPN) of sulfur-oxidizing and sulfate-reducing bacteria based on culture-dependent data. Our taxonomic profiling, using both shotgun and amplicon data, revealed a high diversity of sulfur cycle bacteria, which was found to be dependent on pH levels. Additionally, our findings confirmed recent modelling of specific taxa biogeographical distribution, such as the sulfur-reducing Mesotoga. Using a functional metagenomics approach, we identified non-canonical taxa involved in dissimilatory sulfur metabolism (e.g., sulfate-reducing acidobacteria and members of the Binatota phylum), and canonical taxa engaged in various oxidative, reductive, and organosulfur transformations (e.g., sulfur-oxidizing alpha-, beta-, and gammaproteobacteria). Furthermore, we discovered that multiple taxa in the studied Technosol encoded different enzymes capable of sulfite transformation and the removal of sulfite from various organosulfonate molecules, thus contributing to the cryptic cycling of sulfur compounds. Estimated MPNs of sulfur-oxidizing bacteria aligned with our shotgun and amplicon data, while those of sulfate-reducing bacteria contradicted functional metagenomic findings. Based on our overall analysis, we support the idea that sulfate-reducers belong to the rare biosphere in soil. We suggest that they behave differently in soils compared to aquatic habitats due to the high taxonomic diversity along with low absolute abundance. Our findings unveil a diverse and unique community of sulfur-metabolizing bacteria that has evolved in soil under severe technogenic pollution, high bulk sulfur content, and fluctuating redox states.

## Introduction

The study of microbiota involved in the sulfur cycle was initiated in the late 19th century through groundbreaking Winogradsky and Beijerinck works (Lens et al., 2002). One of the most well-known experiments that paved the way to study sulfur cycle microbiota (SM, hereafter) was the establishment of the Winogradsky column. The column represents a model of submerged ecosystem with C and S sources. Over time, within the column exposed to light, the microbial community stratifies along oxygen and iron sulfide gradients (Zavarzin, 2006), giving rise to a diverse sulfur-transforming community. What thus initially began as the first observation of microbes directly linked to their ecological traits has now revealed an enigmatic picture of an immensely diverse group of microbes. These microbes are ubiquitous and possess the ability to mediate numerous traceable and cryptic biogeochemical transformations, making it increasingly challenging to assign them to specific processes, transformations, or econiches. Most of SM diversity concentrates within dissimilatory pathways, where inorganic sulfur compounds are metabolized and/or deposited outside the cell (Anantharaman et al., 2018). At least five compounds between the most oxidized (sulfate) and the most reduced (sulfide) sulfur state – elemental sulfur, polysulfides, thiosulfate, tetrathionate, and sulfite – can be oxidized, reduced or disproportionated by at least one distinct group of microorganisms, with many overlapping groups also capable of these processes. Recently, microbes liberating and reducing sulfite moiety from organosulfonates were also recognized in the context of dissimilatory sulfur metabolism (Hausmann et al., 2018; Peck et al., 2019; Ye et al., 2023). The diversity of taxa oxidizing sulfide or reducing sulfate is even bigger. Two major sulfur-related operons, Sox and Dsr gene clusters, can act in a reverse way, thus widening our perspective on a possible genetic diversity and capabilities of the SM. Many sulfur oxidizing bacteria (SOB) demonstrate conspicuous morphology, from well-known sheathed filamentous *Beggiatoa* and *Thiothrix* to recently described giant *Thiomargarita* species (Volland et al., 2022; Ravin et al., 2022). On the other hand, sulfate-reducers (SRB) are unprecedentedly diversified taxonomically, with evaluated number of unique operational taxonomic units (OTU) exceeding tens of thousands (Vigneron et al., 2018). In some cases, SRB exhibits a unique natural behavior: they do not proliferate and maintain a low-density population, yet they remain transcriptomically and metabolically active (Hausmann et al., 2019). In that state, cells actively reduce sulfates and significantly influence their ecosystem, which led to the idea of SRB comprising rare biosphere. It is also has become evident that other microbial taxa, not affiliated with SM, can bear complete Dsr operons (Anantharaman et al., 2018). For example, a unique type of metabolism – degradation of complex plant biopolymers coupled to dissimilatory sulfate reduction – was recently discovered in Acidobacteriota members (Dyksma and Pester, 2023). Finally, a rare case of prokaryotic high-taxonomic level endemism has recently been observed in a relic Antarctic lakes. In this habitat, a novel class of bacteria within candidate Electryoneota phylum exists, possessing the complete genetic machinery necessary for both sulfate and sulfur reduction to sulfide (Vigneron, Vincent and Lovejoy, 2023).

SRB and SOB actively participate in elemental cycling beyond sulfur. SRB can degrade hydrocarbons, reduce and precipitate heavy metals from iron to plutonium (Merino et al., 2023), compete with methanogens for substrate (Hu et al, 2022), degrade end- or byproducts of others’ bacteria metabolism; besides, they are also capable of direct extracellular electron transport (Lovley and Holmes, 2022). SOB members can fix CO_2_, oxidize heavy metals, and convert insoluble sulfide minerals. Some phototrophic members of SOB utilize near-infrared light, thus introducing new pathways for energy to enter biogeochemical systems (Beatty et al., 2005). In addition, some SRB and SOB can respire nitrites. Most of SM belongs to microbial dark matter or resists growing in the lab. Many SOB, although have been studied and sequenced using naturally occurring biomass, has yet to be isolated in axenic cultures, while the hundreds of SRB groups are known solely from metagenome binning data (Li et al., 2022; Vigneron, Vincent and Lovejoy, 2023). SM comprise functionally and taxonomically diverse groups of prokaryotes and are crucial for the global ecosystem. However, the modern picture of SM biology and ecology is biased towards the processes in aquatic environments (Li et al., 2022) (oceans, lakes, groundwater) and controlled biotechnological systems (bioleaching tanks, corroded metal structures). Among terrestrial habitats, rice paddy fields, peats and volcanic muds are well represented in current studies, while a wide spectrum of soils, phyllosphere, arid environments, caves and host-associated consortia are largely overlooked. Furthermore, for almost a century, we have been witnessing the emergence of entirely new types of environments because of human activities and the interactions between technology and nature. These include highly contaminated Technosols, landfills, urban areas, as well as metal and/or concrete infrastructure. All these niches are readily colonized by SRB and SOB.

Because of the development and active implementation of functional metagenomic approaches, it has become easier to characterize complex microbial consortia. The binning and subsequent annotation of MAGs allows not only for comprehensive studying of new species, but also hints on the possible ways of their isolation in the laboratory culture. However, metagenomic methods alone are unable to resolve complexity of an environmental microbial consortia; it is important to combine culture-dependent and molecular techniques. In the present work, we describe the soil microbiota of a unique ecosystem formed at a site of a dried-up lake, characterized by severe polymetallic, bulk sulfur, and PAHs contamination.

## Materials and methods

### Design of the study

The study aimed to characterize the sulfur cycle microbiota in the polluted soils of the former Atamanskoe Lake. The first step involved analysis amplicon sequencing data covering the entire lake territory and adjacent soils. In total, 27 samples were taken, with 22 belonging to the lake territory and 5 to the adjacent areas. These samples were used for DNA extraction and amplification of the 16S rRNA gene. Based on the analysis of that data, 9 samples were selected for 1) determination of the physicochemical properties of the soils, 2) total DNA extraction and sequencing, 3) enumeration of sulfate-reducing and sulfur-oxidizing bacteria by MPN approach.

### Object description and sample collection

The soils of the dried-up Atamanskoe Lake, located at the South of Russia, served as the object of study. The lake was initially formed as an oxbow of a nearby river. As a result of the usage of the lake’s territory as a sludge collector from the 1960s to the mid-1990s, a large part of industrial waste from chemical production accumulated in its bottom sediments. Over the past two decades, a Technosol has formed based on the sediments, which is characterized by unprecedented levels of PAH and HM concentrations (Linnik et al., 2022). The amount of technogenic sludge in the lake is about 400,000-420,000 m^3^, containing up to 98,000 mg kg^-1^ of zinc, 1,050 mg kg^-1^ of lead, 13.6 mg kg^-1^ of mercury, 52 mg kg^-1^ of arsenic, and 303 mg kg^-1^ of copper (Minkina et al., 2019). Compared to average lithospheric values, much higher concentrations of Zn, Cd, Pb, Cu, Ni, and Cr are observed, hundreds of times higher for Zn, ten times higher for Cd and Pb, and several times higher for Cu, Ni, and Cr (Bauer et al., 2018). The total concentration of PAH ranges from 499 to 7,178 mg kg^-1^ (Sushkova et al., 2020). According to the World Reference Base for Soil Resources (WRB), the soils are classified as Spolic Technosols. Over half of the minerals in Atamanskoe soils are sulfate minerals, and a sulfide geochemical barrier operates in the deep soil layers (Bauer et al., 2018). As a control, Fluvisol soil from a floodplain site near Lake Atamanskoe with background values of PAH and TM indicators was taken. Soils were collected by taking an average sample from five points, packaged in sterile plastic bags, and stored at t=4°C until subsequent analysis (ISO 10381-1, 2002). For the screening amplicon study, 27 samples were selected, of which 22 represented the lake area and 5 were background soil samples. For the determination of physicochemical properties, microbiological cultures, and shotgun metagenomic analysis, 9 samples were chosen from the 27. The map of soil sample collection is presented in Figure 1.

**Figure 1.**
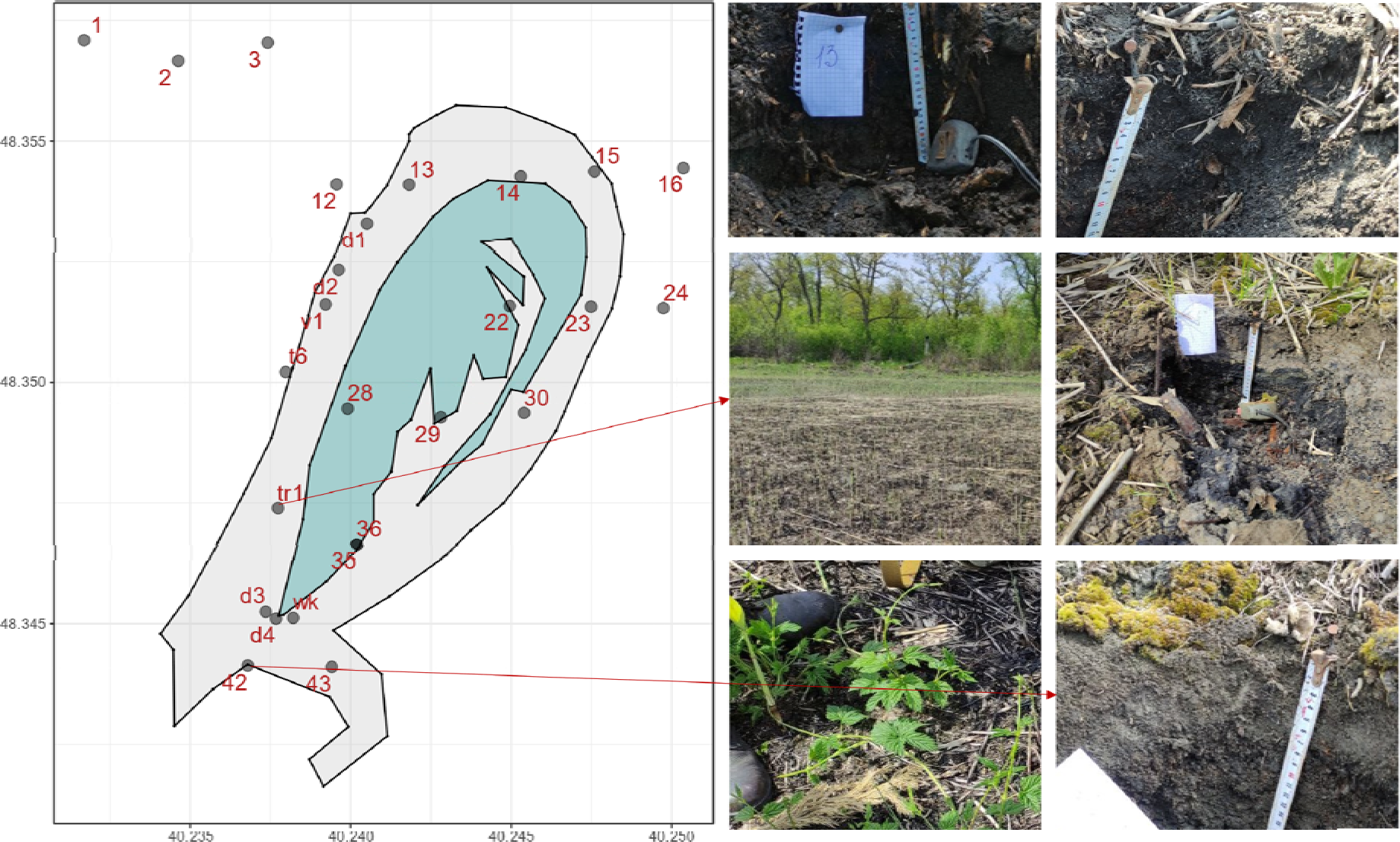
Soil sample collection map: the green zone indicates a former island with abundant vegetation cover, the gray zone represents the area of the Technosol (former water area).

### Soil physicochemical analysis

The data on pH, concentration of PAH and HM mentioned in the study were obtained using the methods specified in (Gorovtsov et al., 2021). In this study, the data were used to analyze their correlation with the content of SM. The table containing the soil parameters and pollutants concentrations data is provided in the supplementary material (file S1, table1).

### DNA extraction and sequencing

#### Metabarcode sequencing

According to the manufacturer’s instructions, total DNA was extracted using the FastDNA SPIN Kit for Soil (MP Biomedicals, UK) for sequencing the 16S rRNA gene. Metagenomic libraries were prepared following the “Preparation of 16S rRNA Gene Amplicons for the Illumina MiSeq System” protocol. The amplification of the V3-V4 region of the 16S rRNA gene was performed using the following prokaryotic primers: forward - TCGTCGGCAGCGTCAGATGTGTATAAGAGACAGCCTACGGGNGGCWGCAG; reverse - GTCTCGTGGGCTCGGAGATGTGTATAAGATACAGGTATCACCHG. Sequencing was carried out on the MiSeq platform (Illumina) with v3 reagents (600 cycles) at the Interdisciplinary Shared Facility of Kazan Federal University (Amplicon P. C. R., Clean-Up P. C. R., Index P. C. R. 16S Metagenomic sequencing library preparation, 2013).

#### Shotgun metagenomics

Total soil DNA was extracted using the FastDNA™ Spin Kit for Soil. For library preparation, the NEBNext Ultra II DNA Library Prep Kit (NEB) was used according to the manufactu er’s instructions. Genomic DNA was fragmented using the Covaris S220. The quality of the prepared libraries was assessed using the 2100 Bioanalyzer (Agilent) with the DNA High Sensitivity Kit (Agilent). The DNA concentration in the samples was quantified using the Qubit 2.0 instrument. Sequencing was performed on the MGI platform (DNBSEQ).

### Bioinformatics and statistical analysis

#### Analysis of 16S rRNA gene pool sequencing data

The filtration of raw sequencing data, quality control, removal of chimeras, and merging of paired reads were performed using Trimmomatic, Pandaseq, and VSEARCH programs. Subsequently, the filtered reads were fed into the Kraken2 program (Wood, Lu & Langmead, 2019) and classified using the SILVA-138 database. The results from Kraken2 were further processed using the Bracken software.

### Analysis of shotgun sequencing data

#### Quality control

Reads corresponding to human DNA were removed using the BMTagger program. Read duplicates were removed using clumpify from the bbmap package. The FastQC program was used for quality control. Adapter sequences, low-quality reads (Q < 10), and short reads (< 40 bp) were removed using Trimmomatic ver. 0.38.

#### Analysis of metagenomic taxonomic profiles

Quality-filtered reads from the quality control step were subjected to taxonomic identification using the Kraken2 program with the PlusPFP dataset. Re-estimation of taxon proportions in the community based on the reads counts was additionally performed using the Braken program. Further analysis and visualization were conducted using R 4.3.2.

#### Assembling and binning

Quality-controlled reads were assembled into contigs using metaSpades (Nurk et al., 2017) default parameters. The contigs were then binned using three independent programs with default parameters: Metabat2, CONCOCT, and Maxbin2. The bins obtained using these tools were merged into consensus bins using the DAS-Tool program (Kang et al., 2019; Alneberg et al., 2013; Wu, Simmons & Singer, 2016; Sieber et al., 2018). To obtain all the reads that were dropped in the process of contigs assembly, these contigs were indexed using bowtie2-build command, and used for raw metagenomic reads mapping (bowtie2 –local --un-conc) saving paired unaligned reads. Further processing of the bins, including contig removal based on tetranucleotide frequency, GC content, and presence/absence of clade-specific marker genes, was performed using the MAGpurify program (Nayfach et al., 2019). Quality control and assembly statistics of the bins were assessed using the CheckM program (Parks et al., 2015) and MiGA online resource (https://disc-genomics.uibk.ac.at/miga/, Rodriguez-R et al., 2018). The bins that passed all quality control steps, had an assembly completeness >60%, and contamination level <15% were considered MAGs.

#### Annotation

MAGs were annotated using the GhostKoala (https://www.kegg.jp/ghostkoala/) and EggNOG (http://eggnog-mapper.embl.de/) online mappers. Based on the annotation outputs, conclusions about reconstructed species in the community were made. All quality-filtered MAGs were taxonomically classified using TypeMat and GTDB datasets available in MiGA. If taxonomic assignments varied between the two datasets, GTDB inference were used.

#### Alignment

Alignments for DsrD and 30S ribosomal subunit protein sequences were performed using ClustalW with slow/accurate setting parameters (https://www.genome.jp/tools-bin/clustalw). The final trees were inferred out of 100 bootstrapped trees using PhyML v20160115.

#### Operon structure determination

In those cases where operon structures are discussed, we manually investigated EggNOG and Operon-mapper (https://biocomputo.ibt.unam.mx/operon_mapper/) outputs to decide whether specific ORFs comprise a unite operon.

#### Metabolic role assignment for a MAG

We used rules developed by (Anantharaman et al., 2018) to decide whether a microbe under a specific MAG contributes to reductive or oxidative processes in sulfur biogeochemical cycling. Species encoding Sat+AprAB+DsrABC+DsrD were considered sulfate-reducers, while those containing DsrAB+DsrEFH were considered sulfur-oxidizers with oxidative-type Dsr pathway.

#### DNA-DNA hybridization

A tool developed by (Meier-Kolthoff et al., 2022) was used to calculate hybridization statistics for metagenome-assembled genomes (https://ggdc.dsmz.de/).

### Quantitative analysis of sulfate-reducing bacteria

Enumeration of sulfate reducers was carried out using the most probable number (MPN) method as follows: Cells were separated from soil particles in a sterile mortar using a rubber pestle (soil-to-water ratio of 1:10), and then shaken on a rotary shaker in flasks for 40 minutes. A 1:10 cell suspension, after separation from soil particles, was serially diluted up to 10^-7^. Each dilution, in 4 replicates, was then used to inoculate a tube containing 20 ml of modified Postgate B medium with pH of 7.5, the composition of which is specified in (Kushkevych, 2020). The tubes were sealed with rubber stoppers and incubated at t=30°C for three weeks, after which the number of replicates showing iron sulfide precipitation was recorded for each dilution. The recorded data were used to calculate the most probable number using with the MPN package in R 4.3.2., which utilizes a modified version of the classical MPN calculation (Ferguson & John, 2019).

### Quantitative analysis of sulfur-oxidizing microorganisms

Soil dilutions were prepared as described above. Then, 1 milliliter of each dilution was used to inoculate 10 mL of Waksman medium with elemental sulfur (pH = 7.5), poured in 50 mL plastic containers 50 mL (each dilution in triplicate). Incubation was carried out for two months, t=25°C. The presence of growth was determined based on three indicators: a decrease in pH to values <5, visible solubilization of elemental sulfur, and visible microbial growth/turbidity. A positive result was considered when the medium exhibited an acidic reaction after the incubation period, and microscopy of solubilized sulfur demonstrated the presence of cells. The medium’s reaction was tested using acidity indicator solutions (methyl red, bromocresol green). MPN was calculated as described above.

### Quantitative analysis of thiosulfate-oxidizing microorganisms

Soil dilutions were prepared as described. Then, 100 microliters of each dilution were used to inoculate Beijerinck medium with thiosulfate and pH indicator bromocresol green (0.04%), distributed at 600 µL per deep-well polystyrene plate with a well volume of 2.5 mL (each dilution in quadruplicate). Incubation was carried out for one month, t=25°C. The presence of growth was determined by a color change of the pH indicator from violet-blue to yellow and the formation of a yellow cell precipitate at the bottom of the well. MPN was calculated as described above.

### Statistical analysis

Standard statistical tests, such as the Shapiro-Wilk test for normality of distribution, the pairwise Wilcoxon test, and Spearman correlation analysis, were carried out using base packages in the R 4.3.0. If it was necessary to compare groups of unequal sample sizes, the required statistical test was performed based on 100 random subsamples of a bigger group, with a size of each subgroup equal to the overall size of a smaller group). Calculation of biodiversity indexes and ecological distances was also performed in R using the vegan package (Oksanen, 2013). Ecological distances were inferred using bray-curtis dissimilarity, significance of NMDS clusters difference was assessed by PERMANOVA (n permutations = 999).

## Results and discussion

### Metabarcode screening shows predominance of spore-forming SRB and acidophilic chemolithoautotrophic SOB in Technosols

In addition to typical members of the reductive and oxidative stages of the sulfur cycle, such as *Desulfovibrio* and *Thiobacillus*, many other microorganisms are involved in sulfur cycling. Therefore, for comprehensive screening of the obtained metagenomic data, keyword searches, systematic reviews and articles containing information about species and genera with documented metabolism of inorganic sulfur were used. Information on the oxidative stage was obtained from studies (Ghosh & Dam, 2009; Kumar et al., 2018), on the reductive stage from studies (Rabus, Hansen & Widdel, 2006; Anantharaman et al., 2018), and on sulfur-disproportionating taxa from the review (Slobodkin & Slobodkina, 2019). Only taxa represented by at least 10 reads of the ribosomal gene were taken into consideration. Additionally, sampling sites were divided into two groups – background (control) soils and Technosol samples – to compare the SM diversity. The screening results are presented in Figure 2. A total of 28 genera associated with dissimilatory metabolism of sulfur compounds were identified in the dataset. The majority of taxa were represented by a small number of the reads. The most abundant were the genera *Desulfosporosinus* (1927 reads), *Desulfallas* (713 reads), *Desulfofarcimen* (593 reads), and *Desulfitomicrobium* (468 reads). Among them, *Desulfosporosinus* surpassed the rest both in terms of read count and its proportion, reaching up to 3-4% of the entire metagenome in some samples (d2, 13, 23, 30). The greatest taxonomic diversity was observed in samples 43 and d3, while the lowest was observed in samples t6, 12, and 3, with no sulfate reducers detected (according to established criteria) in samples 16, 28, 24, 2, 1, 36 at all. Notably, there is a predominance of spore-forming genera belonging to the Bacillota phylum: *Desulfosporosinus*, *Desulfofarcimen*, *Desulfitomicrobium*, *Desulfurispora*, *Thermincola*, and *Pelotomaculum*. This may reflect the highly changeable conditions of Atamanskoe soils, particularly the fluctuating redox potential due to the periodic flooding.

**Figure 2.**
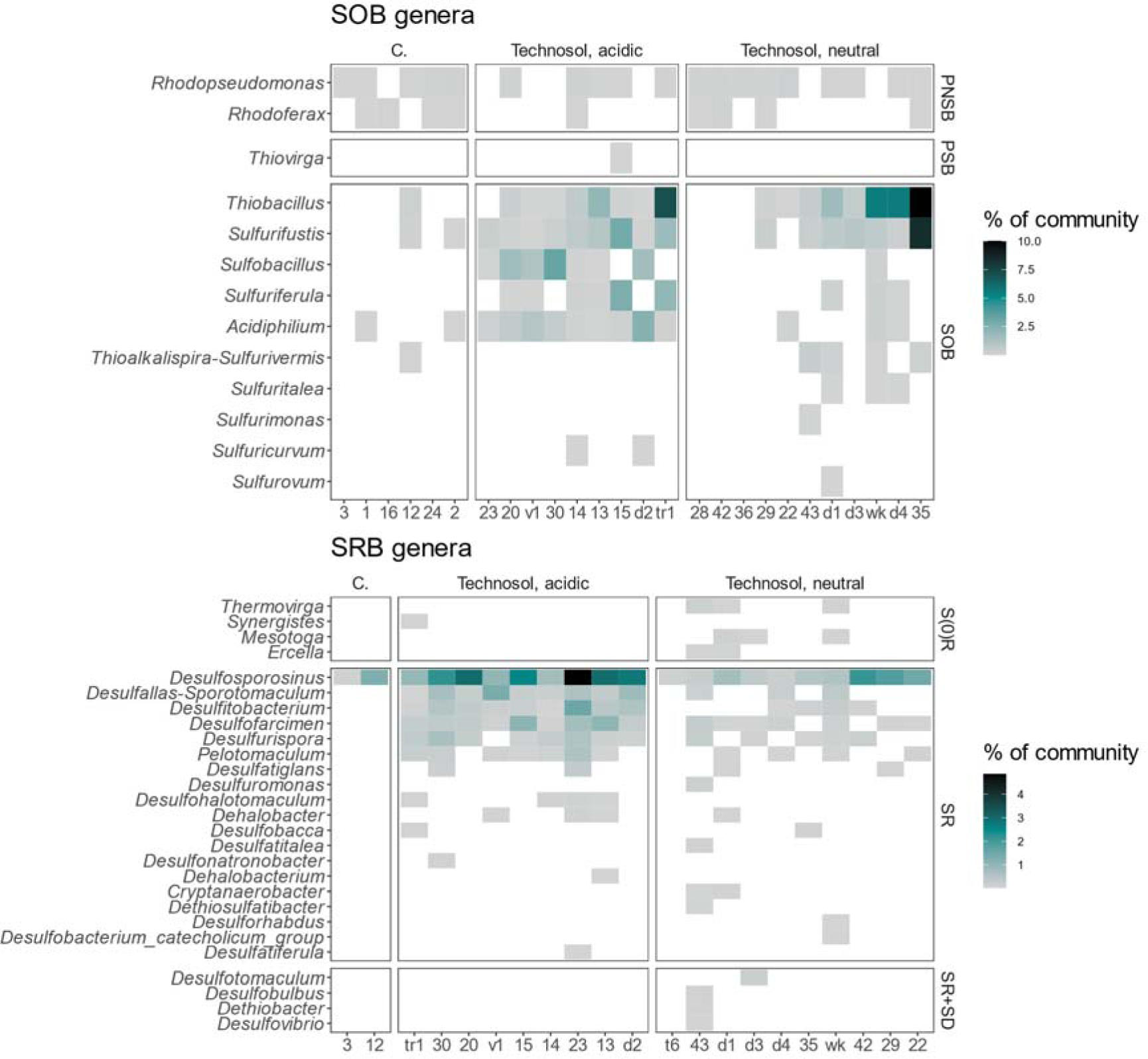
Profile of SRB and SOB genera based on amplicon data; SR - sulfate reducers, SD – disproportionating microorganisms, S(0)R – sulfur-reducing bacteria; SOB – autotrophic sulfur oxidizers; GSB - green sulfur bacteria; PSB - purple sulfur bacteria; PNSB - purple non-sulfur bacteria; C. – background soils.

Among the sulfur bacteria, 13 genera were identified, of which 2 were phototrophs. No significant diversity differences or dominance within or between samples were observed for the PNSB group. A small number of reads were recorded for the purple sulfur bacteria *Thiovirga*. Many colorless sulfur bacteria were recorded. *Thiobacillus*, *Sulfobacillus*, *Sulfurifustis Acidiphilium*, and *Sulfuriferula* were the predominant genera. The most diverse sites were 43, 35, d1, d4, d3, and 14. *Thiobacillus* reached exceptional proportions of reads in samples d4, wk, tr1, and 35, accounting for 5.5%, 5.4%, 7%, and 10% of all sample reads, respectively. The purple non-sulfur bacteria were distributed most evenly among the groups, although overall diversity was higher in Atamanskoe samples. In the control soils adjacent to the lake area, only a small number of sulfur-oxidizing taxa were detected. Figure 3 represents the spatial distribution of the sulfur microbiome across the Atamanskoe territory.

**Figure 3.**
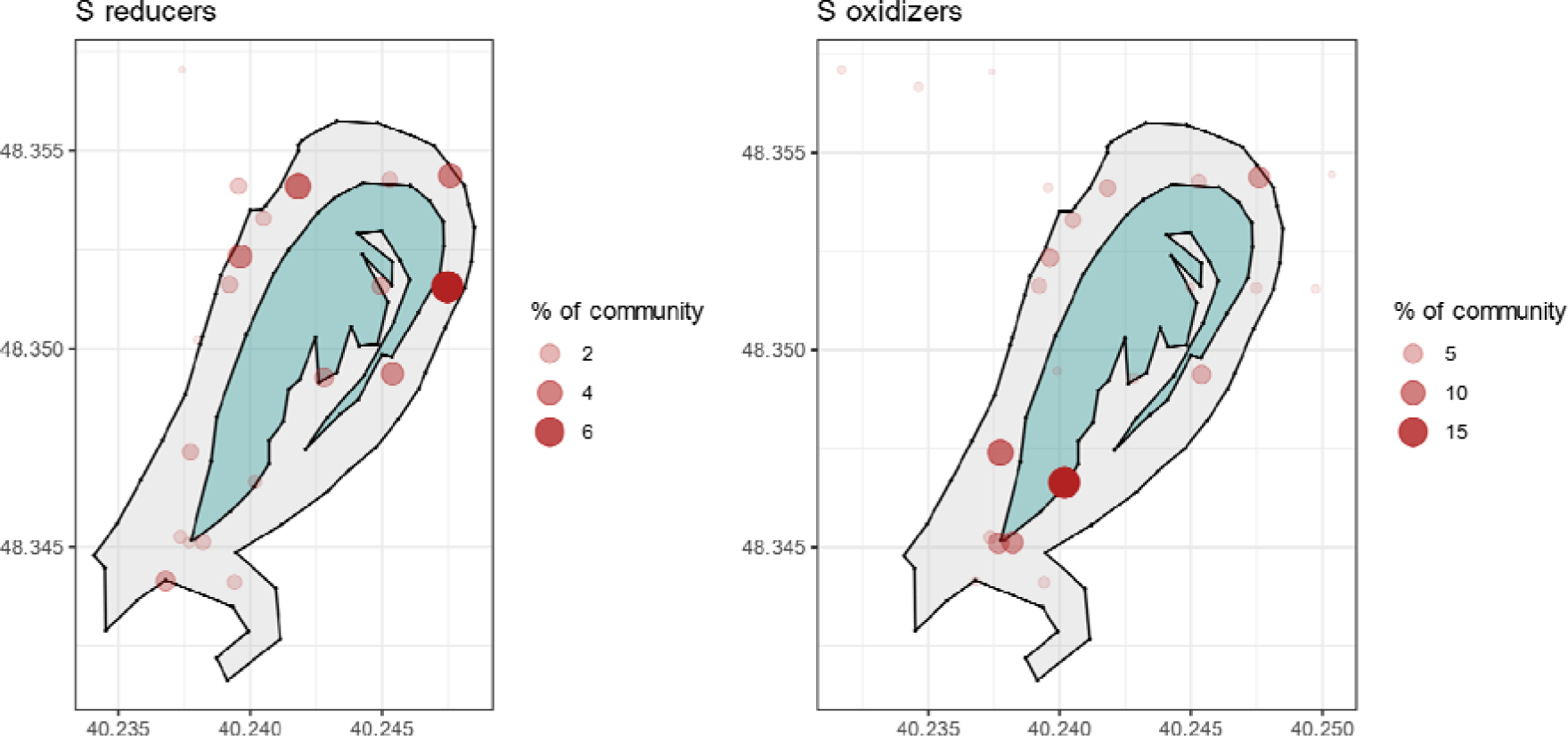
Spatial distribution of studied sulfur microbiome. Each point represents the sum of bacteria with reductive or oxidative metabolism correspondingly in the sample.

Based on the above data, it can be concluded that there is a high heterogeneity in the sulfur microbiome of the soils formed in the lake territory. It is worth noting that for such non-systematic and diverse groups of bacteria like sulfate reducers, the use of ribosomal genes may not be informative for community classification. False-positive signals and an inflated number of unique genera can be expected. However, all of this indicates a greater diversity of 16S rRNA gene variants. Additionally, PCR errors and variations in primer annealing can significantly distort the community profile. For a more in-depth study of the microbial diversity in the sulfur cycle, we selected nine samples for shotgun sequencing and further investigation.

### Analysis of total soil DNA shows predominance of SM, but contradicts taxonomy distribution inferred from amplicon data

Before assembling reads into contigs, the workflow for sequencing total soil DNA was not different from that for amplicons. The same taxa were selected for analysis as in the amplicon metagenomes. A minimum threshold of 1,000 reads was set for including a genus in the analysis. Figure 4 presents the profile of sulfate-reducing bacteria in 9 soil samples. The abundance and diversity differ significantly from that of the amplicon profile. *Desulfosporosinus*, *Desulfitobacterium*, and *Desulfofarcimen* are no longer dominant. Among dominant SRB were *Desulfuromonas*, *Pseudodesulfovibrio*, *Desulfovibrio*, and *Mesotoga*; these genera reached >0.1% of the community in case of several samples, which is typically unusual observation for sulfate-reducers in soils. *Desulfuromonas* and *Pseudodesulfovibrio* reached comparable abundance in background soil. Many more minor taxa were either abundant in Atamanskoe metagenomes or absent in background soil, such as *Petrimonas*, *Desulfoglaeba*, *Desulfobulbus* and *Desulfomicrobium*. An interesting case represents *Mesotoga*: the analysis identified comparatively high read numbers for this genus (14541 in d3 sample). *Mesotoga* bacteria are unique Thermotogota members thriving in mesophilic conditions. It was previously suggested that anthropogenic activities, especially associated with PAH contamination, favor *Mesotoga* distribution across different environments (Nesbø et al., 2019). Moreover, these microorganisms can promote growth of hydrogenotrophic sulfate-reducers (Fadhlaoui et al., 2018). Along with *Mesotoga*, *Desulfuromonas*, *Oceanithermus* and *Campylobacter* represent a significant part of the Atamanskoe SRB community capable of sulfate and sulfur reduction or sulfur reduction solely.

**Figure 4.**
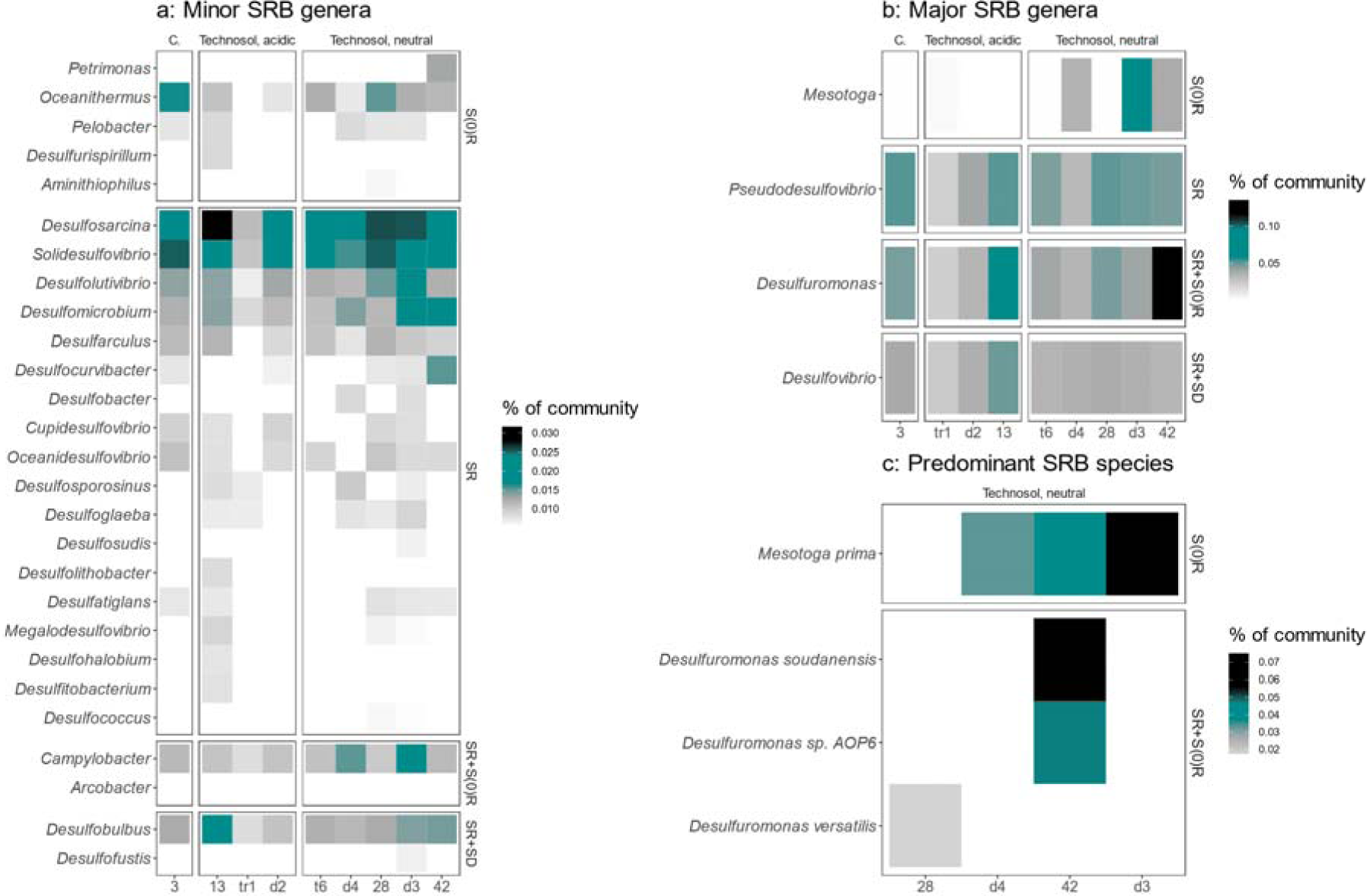
Profile of SRB genera based on total soil DNA sequencing; SR - sulfate reducers, SD – disproportionating microorganisms, S(0)R – sulfur-reducing bacteria; C. – background soils.

Sulfur-oxidizing taxa identified by shotgun sequencing are presented in Figure 5. *Sulfuriferula*, *Sulfurifustis*, *Sulfuricaulis*, and *Thiobacillus* are present in high numbers in the tr1, 42 and d4 sites. The most abundant of all SOB was *Thiobacillus* genus, reaching almost 10% of all tr1 sample and 5% of d4 sample. These samples also have demonstrated the highest MPN values for sulfur and thiosulfate oxidizing microorganisms (see below). The second was *Sulfuriferula* (almost 2% and 336626 total reads of tr1 sample). Within it, *S*. *plumbiphila* was predominant, representing, as was in the case of *Mesotoga*, the technogenic nature of the studied ecosystem. The same pattern can be observed for the other samples and groups: higher abundance and diversity of SOB in Atamanskoe soils. Moreover, several identified taxa are known to be electroactive, such as *Rodoferax*, *Thioclava*, *Acidithiobacillus* and *Desulfuromonas*. In general, 45 and 76 unique genera associated with sulfur cycle were identified in Atamanskoe soils based on amplicon and shotgun data respectively.

**Figure 5.**
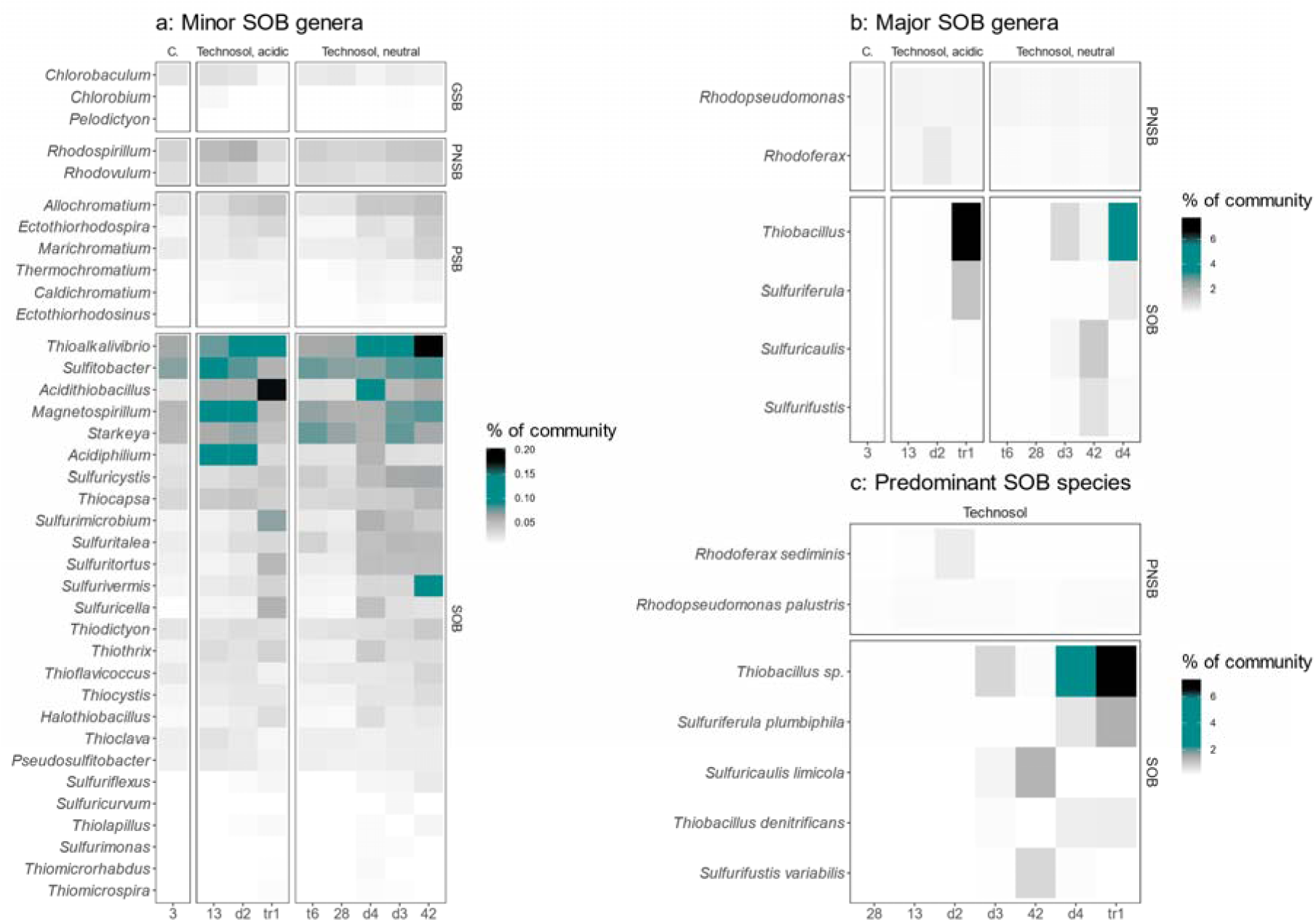
Profile of SOB genera based on total soil DNA sequencing; SOB – autotrophic sulfur oxidizers; GSB - green sulfur bacteria; PSB - purple sulfur bacteria; PNSB - purple non-sulfur bacteria; C. – background soils.

### Pollutants content is associated with SM abundance

In addition to acidity level, the content of PAHs and certain HMs was also significantly associated with both the abundance and diversity of the sulfur cycle microbiota. Table 1 presents the results of the Spearman correlation analysis between these indicators and the concentrations of Cu, Pb, Zn, Cd, and the sum of 16 types of PAHs.

**Table 1.** Spearman correlation coefficients between pollutant concentrations and statistics of sulfur microbiome. ‘ns*’* – non-significant, ‘*\*’* – *p* value < 0.05, ‘*\*\*’* – *p* value < 0.01.

| Parameter | Zn | Pb | Cd | Cu |
| --- | --- | --- | --- | --- |
| SRB abundance | ns | ns | 0,5 ** | ns |
| SOB abundance | ns | 0,41 * | 0,54 ** | ns |
| SM abundance | ns | 0,39 * | 0,55 ** | ns |

The diversity of total SM abundance was significantly correlated with bulk Cd content, as both SRB and SOB demonstrated moderate correlation with this indicator, while Pb were significantly correlated only with SOB. It is worth noting that Zn is the main pollutant in Atamanskoe soils. Zn concentration averages 3,3% in studied Atamanskoe soils (supplementary file S1, table 1), with the maximum value of 72886 µg kg-1 in sample t6. The maximum recorded concentration of bulk Zn content for the whole period of ecological monitoring in Atamasnkoe soils was up to 15% (unpublished data). Such concentrations are typically observed for Zn ores (Wang, 2016). However, no significant correlations were found between sulfur microbiota abundance and Zn content. The tolerance to extreme Zn content in polluted areas is documented for microbial species, especially for acidophilic and/or sulfur-metabolizing bacteria (Watkin et al., 2009; Mangold et al., 2013), and for multiple photosynthetic bacteria including purple non-sulfur genera such as *Rhodopseudomonas* (Edwards et al., 2013), which detoxify Zn by its metal sulfide precipitation. As for Cd correlations, it is also known to be associated with sulfur cycle microbiota (Naz et al., 2005; Zheng et al., 2015; Ramos-Zúñiga et al., 2019; Li et al., 2019; Zheng, Wu & Sun, 2021; Yin et al., 2024). Recent study has shown increased cadmium tolerance and sulfur oxidation rates in microcosms stressed with high sodium sulfate concentrations (Wang et al., 2021).

These results show that overall SM content response differently to different heavy metals, and that even metal ore-comparable concentrations of some metals can be positively correlated with SM in soils.

### Diversity of sulfur microbiota differs from whole metagenome diversity and depends on pH

We conducted an ecological analysis using two approaches. Firstly, we assessed the alpha- and beta-diversity of the Atamanskoe soil microbial community by considering all identified genera (whole-metagenome diversity). Secondly, we focused on the sulfur-metabolizing community as a separate group and re-evaluated diversity measures specifically for this group (sulfur microbiota diversity). Our findings highlight pH as the primary driver shaping the sulfur microbiota (Figure 6).

**Figure 6.**
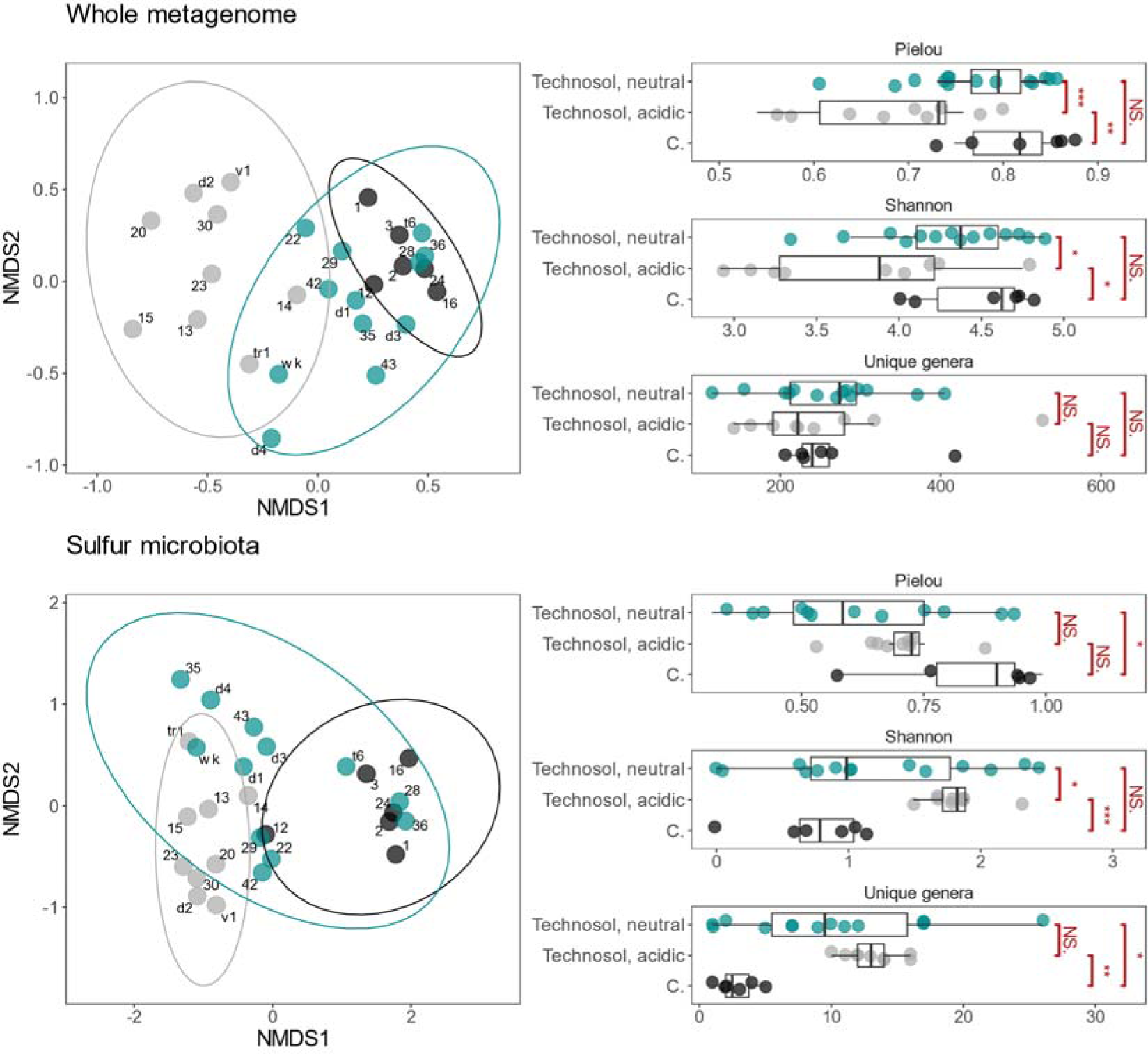
Analysis of alpha- and beta-diversity of the studied soils based on the content of all genera (top) and sulfur cycle microorganisms (bottom); ‘*’ = *p*-value < 0.05, ‘**’ = *p*-value < 0.01, ‘***’ = *p*-value < 0.001, ‘NS.’ = non-significant.

In neutral soils, both the whole-metagenome diversity and sulfur microbiota diversity exhibited high heterogeneity, encompassing both poor and diverse microbial communities at the same time. However, in acidic soils (pH < 4.5), although the whole-metagenome diversity remained similar to the neutral group, the sulfur microbial community displayed higher diversity along with low data variance. These soils harbored a higher abundance of sulfur-related taxa compared to the neutral soils, and the mean Shannon diversity was significantly higher in the low pH group (*p*-value < 0.05). Interestingly, the whole-metagenome diversity showed the opposite trend: acidic soils generally exhibited lower richness in terms of unique taxa and Shannon diversity. Furthermore, beta-diversity analysis revealed significant differences in the structure of the sulfur-metabolizing community among acidic soils, neutral soils, and the control group. Although a similar effect could be observed for the whole-metagenome community structure, it was less pronounced, with neutral Technosol samples clustering closer to the control samples. It is also interesting to note that, despite severe Zn (and other metals) pollution, diversity levels in Atamanskoe were comparable to that of control soils. It may be suggested that even under such pollution levels the precipitation of HM as metal sulfides by sulfur cycling bacteria is beneficial for other members of soil microorganisms and helps maintain the sustainable and diverse microbial community.

### Enumeration of culturable SRB and SOB demonstrates differences in their ecology

Both SOB and SRB groups are hard to culture and are based on physiological rather than taxonomic criteria, i.e. the ability to oxidize or reduce sulfur compounds respectively. For such groups of microorganisms, calculation of the most probable number as a measure of their abundance is a popular approach. An alternative is to use qPCR for Sox or DsrAB gene copies enumeration. Still, there is no universal and at the same time accurate technique for the counting of these groups, and all methods have their own constraints. We aimed to estimate SRB and SOB using the common approach, that is, MPN, and compare possible inferences with those made using metagenomic data. MPN calculation is still extensively implemented in oil and gas industry (Shen and Voordouw, 2017), as well as in geochemical and metal corrosion studies, and, in the case of SRB, demonstrates positive correlations with qPCR approach (Abdulina et al., 2020; Zambrano-Romero et al., 2023). Enumeration of SRB was conducted during the spring and autumn seasons over a period of two years, from 2021 to 2023, while the seasonal variation in the abundance of sulfur-oxidizing microorganisms was not considered and was assessed only during spring 2023. (Figure 7).

**Figure 7.**
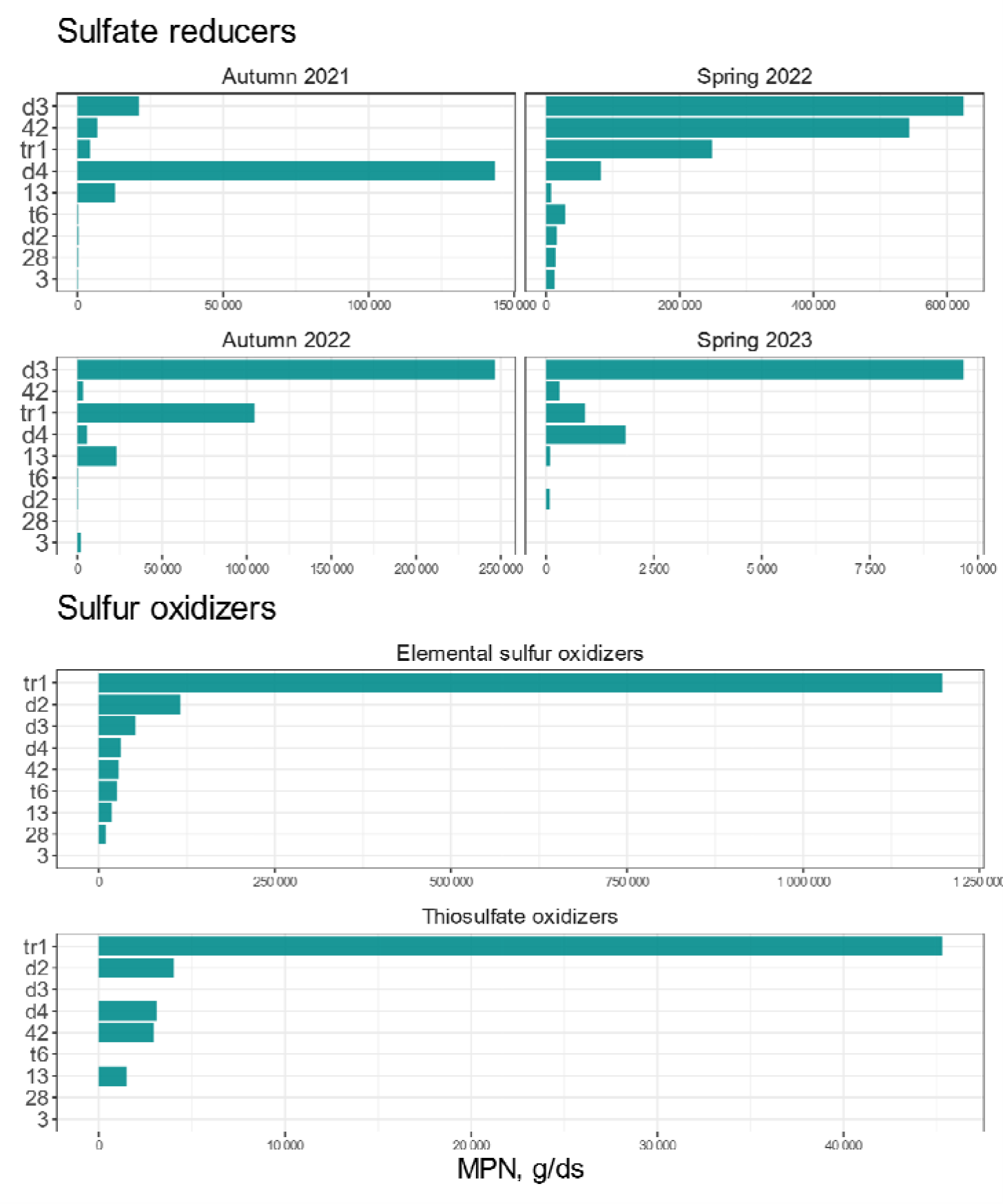
Quantitative assessment of SRB and SOB (most probable number of cells in one gram of dry soil material)

Estimates for SRB ranged from hundreds of cells per gram of dry soil (c gds^-1^) to hundreds of thousands in different seasons and years between samples. The highest value was recorded in sample d3 during spring 2022, reaching over 6×10^5^ c gds^-1^., while the lowest value was observed in sample 28 during autumn 2022, with less than a hundred c gds^-1^. Seasonal dynamics revealed both oscillations of SRB quantity in studied soils and its predominance in Technosol samples. The absolute numbers of SRB cells in Atamasnkoe were similar to that of typical numbers observed in certain groups of soil microorganisms. For instance, soil yeasts typically range between thousands and hundreds of thousands c gds^-1^, while culturable actinomycetes can reach counts in the millions c gds^-1^ (Pulikova et al., 2022). Estimated by the culture-dependent method SRB content demonstrates only a tendency rather than strict correlation: samples with high quantity of SRB-affiliated reads show highest MPN values. One should also consider that laboratory cultivation conditions (such as permanent anoxia, excess sulfate concentrations and readily available nutrients) are not the same as in soil, which may “smooth out” the true representation of the active portion of soil SRB population. Moreover, a typical approach is to express SRB community in cells counts or Dsr gene copies per volume or a mass of specific material. It might be correct in case of a water ecosystem, since oxygen gradient is practically vertical and sulfate content is high, which leads to more or less linear relationship between SRB content physico-chemical parameters. Soils, though, are highly heterogeneous. Redox changes, particle-size distribution and availability of multiple, often solid-state electron acceptors dramatically influence the distribution of this, mostly anaerobic, group of bacteria. Obtained MPN values indicate that SRB in soils indeed can be a part of rare biosphere and exhibit different behavior from that of aquatic habitats.

Abundance of sulfur compounds oxidizers varied much less compared to SRB, reaching tens of thousands c gds^-1^ for both groups in all samples, except for tr1, where the minimum and maximum values were 4.5×10^4^ c gds^-1^ for thiosulfate oxidizers and 1.1×10^6^ c gds^-1^ for sulfur oxidizers, respectively. In contrast to the oxidation of elemental sulfur which was observed in all samples, no thiosulfate oxidation was detected during the monitoring period in samples 3, d3, t6, and 28. It is worth noting that media composition and laboratory conditions in SOB enumeration studies typically promote the growth of *Thiobacillus* species (Sattley & Madigan, 2006), however, in this study, no typical *Thiobacillus* cells were observed in any samples with elemental sulfur. Instead, visible filamentous microorganisms with complex morphology were observed throughout. Any occurrence of such microorganisms was accompanied by decrease in pH value by >2 points and visible sulfur solubilization. Interestingly, both amplicon and shotgun results identified *Sulfurifustis* among dominant SOB genera, which is filamentous and can oxidize a variety of sulfur compounds. Universal medium for SOB does not exist, and each set of conditions will favor a specific group or taxon over the others. Moreover, we did not estimate cell numbers of photosynthetic SOB, so reported values should be considered underestimations. However, these results agree well with metagenomic results: SOB are large and impactful part of Atamanskoe’ soil microbial communities.

### Metagenome-assembled genomes

After quality control, shotgun metagenome reads were assembled into contigs, which were grouped into bins using three independent binning programs. A total of 288 MAGs were obtained with assembly completeness above 60% and contamination below 15%. Among them, 146 genomes achieved assembly completeness of >90%. Assembly quality and statistics were evaluated using both MiGA and CheckM software. All the genomes were then annotated using KEGG GhostCoala tool and EggNOG web resource. Raw data produced by EggNOG were searched against a manually compiled list of 104 sulfur cycle genes (supplementary file S1, table 2). A total of 50 MAGs were selected based on the presence of individual genes or gene groups associated with sulfur cycle metabolic pathways. Among them, 24 MAGs reached >90% level of completeness and <5% level of contamination. Taxonomy and assembly statistics for selected genomes are presented in supplementary material (S1, table 3).

### Taxonomy of binned population is highly diverse

Among 50 MAGs, only two were classified at the species level: *Pseudomonas yamanorum* (MAG 42_83) and *Sulfuriferula multivorans* (MAG d4_19). Most of the other MAGs reached only class or order level. Genomes were distributed among the following unique phyla: Pseudomonadota (32), Acidobacteriota (8), Binatota (7), Gemmatimonadota (1), Nitrospirota (1), Chloroflexota (1). Pairwise AAI comparison indicated low taxonomic similarity between 50 assembled MAGs. Only three pairs showed high AAI numbers and were further confirmed to be identical at the species level using *in situ* DNA-DNA hybridization. MAGs tr1_38 and 13_92 are the same species of Binatota phylum (*p* = 97.45%), MAGs d4_45 and tr1_93 – a species within Burkholderiales order (*p* = 97.89%), d4_9 and 42_11 represent a species of Acidiferrobacterales order (*p* = 98.05%). The phylogenetic tree of the fifty bins was constructed based on the multiple alignment of 30S ribosomal subunit sequences (Figure 8).

**Figure 8.**
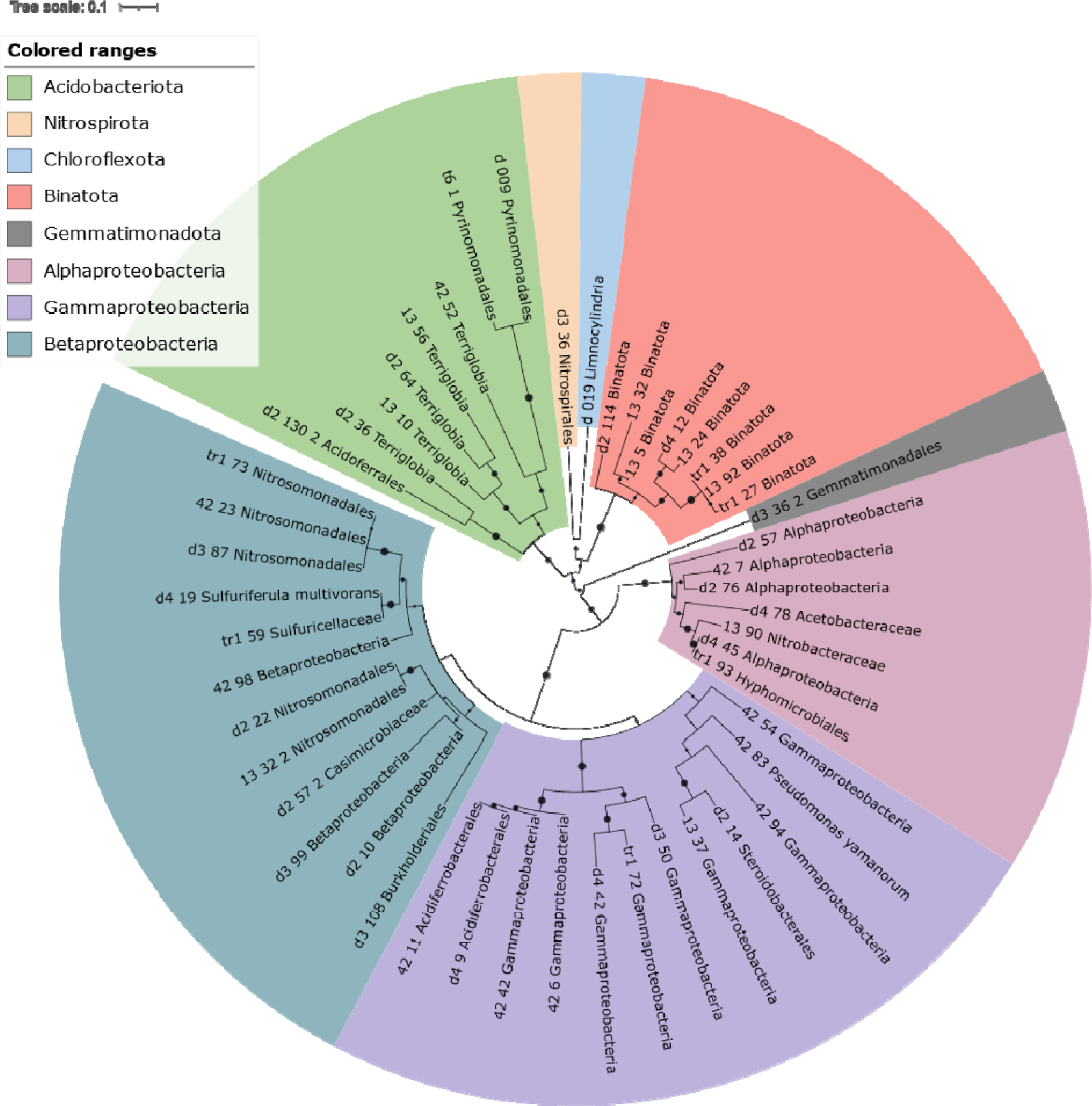
Phylogenetic tree of all assembled MAGs related to the sulfur cycle in Atamanskoe Technosols. Tree was inferred from 30S subunit multiple alignment using ClustalW with the following parameters: PhyML v20160115 ran with following model and parameters: --pinv e --alpha e --nclasses 4 -o tlr -f m --bootstrap 100. Dots in each node represent bootstrap support.

### There were no typical sulfate-reducers among obtained MAGs

The first striking observation was that no MAGs belonging to a typical member of SRB (such as *Desulfosporosinus*, *Desulfovibrio* or *Desulfuromonas*) were assembled. We hypothesized that most of the SRB-affiliated reads have not been used for contigs and bins assembly due to the high taxonomic diversity along with low absolute abundance. To check whether it is true or not, raw reads of soil metagenomes were mapped onto corresponding contigs, and unmapped reads were used to assemble new sets of contigs. The latter were then inspected for the presence of SRB. In most samples, quantity of dropped reads affiliated with SRB were higher or almost identical to that of assembled into initial contigs (supplementary file S2, figure 1). We followed the idea that genomes of diverse but nonabundant species have less chances to be assembled in a typical shotgun metagenome analysis pipeline, and so decided to concatenate all contigs assembled from dropped reads and perform binning on that aggregated dataset. Such an approach yielded an additional group of 37 bins, of which none was associated with known SRB-containing taxa and only two passed quality control. They were added to the main MAGs dataset: MAG d_009 was classified as member of Pyrinomonadales order (Acidobacteriota) and MAG d_019 was the only bin affiliated with Chloroflexota phylum obtained in this study. Other MAGs were investigated for the presence of dissimilatory sulfate reduction genes. Identified genes are shown in figure 9.

**Figure 9.**
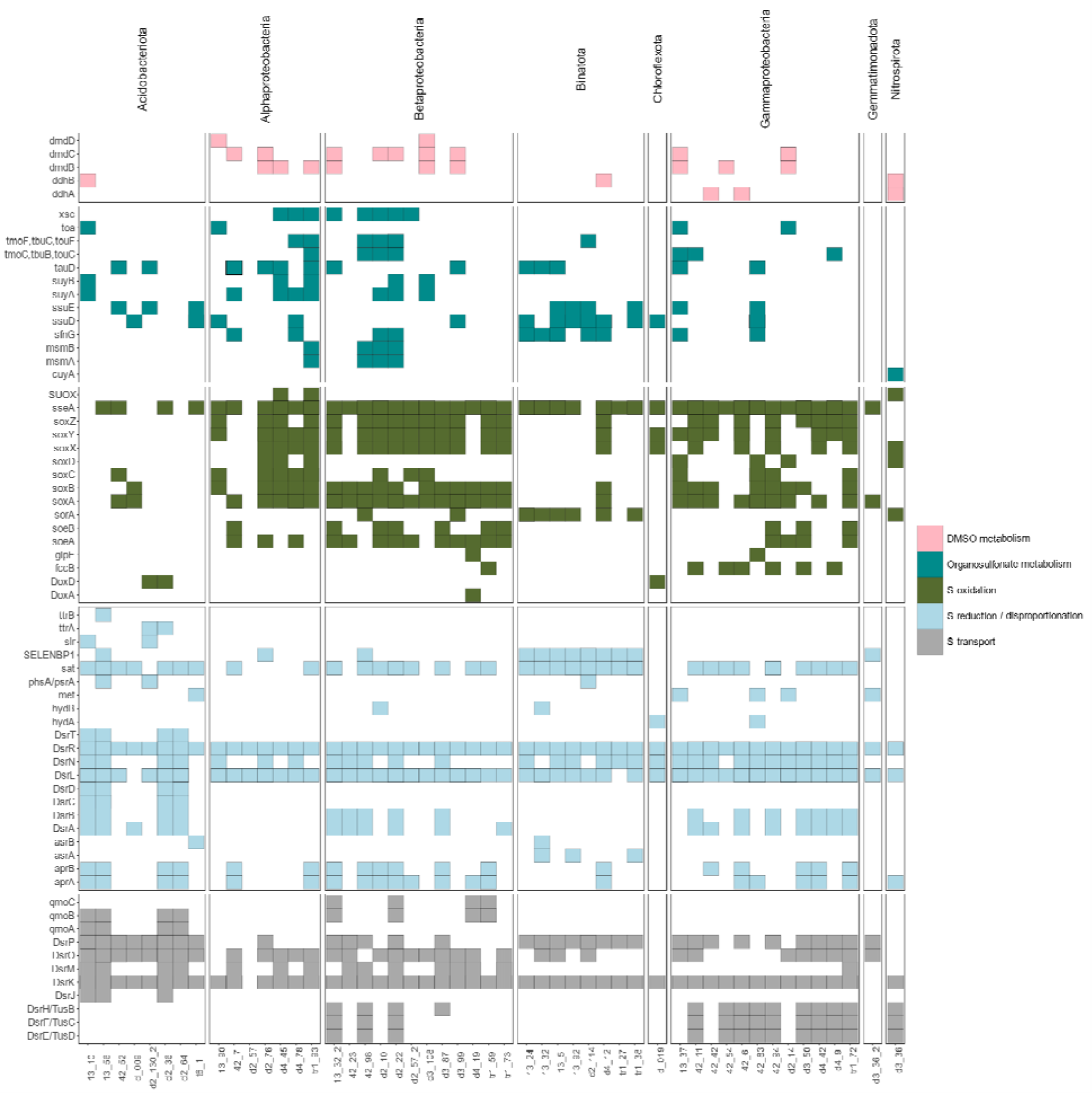
The diversity of sulfur cycle genes and corresponding pathways in all assembled MAGs.

### Binatota members are important part of Atamanskoe microbial community

Seven MAGs were affiliated within Binatota, a diverse and enigmatic candidate phylum. The closest taxon in terms of AAI was *Desulfacinum infernum* for all Binatota MAGs. Although they do not contain Dsr genes, all of them have SELENEBP1 gene, encoding methanethiol oxidase which converts methanethiol to H_2_S. Three of them also contain 1/3 or 2/3 genes that encode assimilatory sulfite reductase (Asr) subunits; six encode different enzymes able to produce sulfite from taurine (tauD), dimethylsulfone (ssuE) and alkanesulfonates (ssuD). Two MAGs have incomplete gene clusters for thiosulfate and sulfur disproportionation (hydB, phsA). Most of the MAGs also encoded dimethylsulfone monooxygenase, an enzyme that produces methanesulfonic acid, which also can be converted to sulfite by toluene monooxygenase tmoF, presented in one of the bins. No AprAB genes were found in this group, but sat gene appeared in all. MAG d4_12 was the only in Binatota group to encode Sox cluster besides the SELENEBP1, ssuD, sfnG and aprAB+sat genes, showing great multifunctionality in the context of sulfur metabolism. Binatota members also seem to be dominant in Atamanskoe soils, since four Binatota MAGs in sample 13 comprise more than 25% of the mapped reads (S1, table 3). Binatota members are widespread, though known only from metagenomic binning data. Recent analysis of Binatota genomes study revealed great metabolic versatility and methylotrophy (including the ability to metabolize sulfomethanes and dimethylsulfone) as a common trait for more the one hundred Binatota-related MAGs (Murphy et al., 2021). It seems that members of the taxa are able to couple methylotrophy to sulfite reduction in studied soils, with dimethylsulfone being central compound in such metabolism, since most of the Binatota bins contained sfnG (dimethylsulfone to methanesulfonic acid) or ssuE (dimethylsulfone to sulfite) or both.

### Acidobacteriota members are predominant sulfate-reducers in studied Technosols

We obtained 8 Acidobacteriota bins, five of which have classified as Terriglobia members, two belong to Pyrinomonadales (Blastocattelia class), and one belongs to a candidate order Acidoferrales. Five Acidobacteriota MAGs encode capability to liberate sulfite from organosulfonates using ssuE, ssuD and tauD, as it was in the case of assembled Binatota MAGs, but also via sulfolactate lyase (suyAB) in the case of 13_10 MAG. The latter also encoded taurine-oxoglutarate transaminase which converts taurine to sulfoacetaldehyde. MAG t6_1 encoded capacity for production of methanethiol from methionine. Acidobacterial bins (d2_35, 13_56, d2_130_2) were the only ones from all dataset to encode ttr cluster genes, such as ttrA and ttrB, which indicates possible tetrathionate reduction. Four MAGs encoded the complete set of genes required for dissimilatory sulfate reduction (sat, AprAB, DsrAB, DsrC) and two encoded 1/3 subunits of Asr and 1/2 of Dsr complex. Recent studies have shown the significance of DsrD gene for assigning DsrAB-encoding bacteria to either dissimilatory sulfate-reducers or sulfur reverse-oxidizers. In our case, EggNOG and KEGG annotator have not been able to show the presence of DsrD, so we manually inspected all the MAGs and have found proteins with putative DsrD function in d2_36, d2_64, 13_10 and 13_56 bins. We used clustalW to perform multiple alignment of these proteins with 18 known acidobacterial-type DsrD genes and 1 Archaeoglobus DsrD variant (as the outgroup), and iTol service to visualize it (Figure 10). These proteins share identical conservative motifs with known acidobacterial DsrD proteins, which means they are highly likely functional in the studied genomes. Visualization of the alignment and corresponding metadata can be found in supplementary file S2, figure 2. Dsr operons in these four genomes demonstrate great variability. We have found some genes to be duplicated, such as DsrN, DsrM and DsrO, and 3/4 bins encoded DsrAB-associated protein of unknown function near the DsrAB (supplementary material file S2, figure 3).

**Figure 10.**
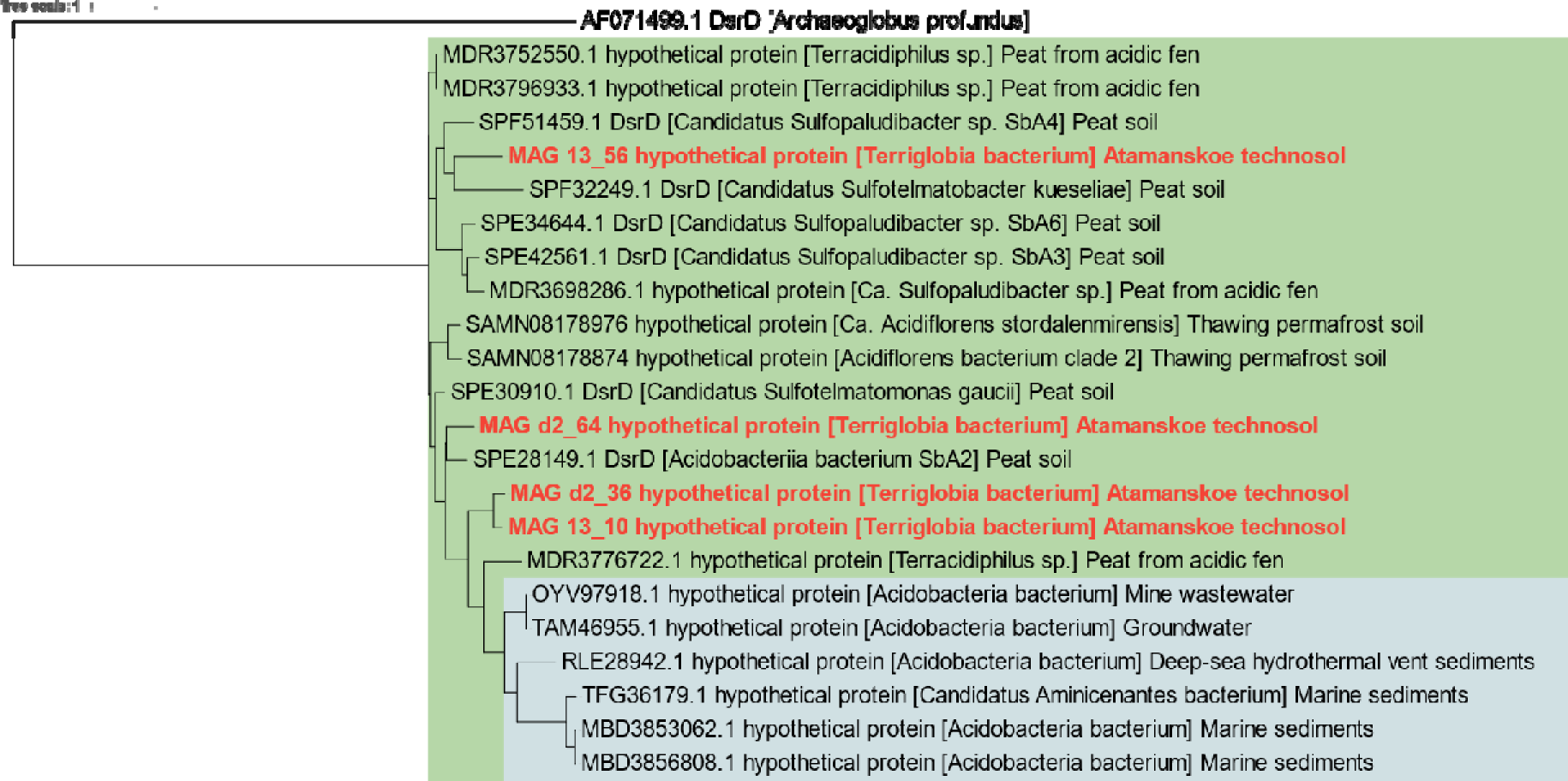
Phylogenetic tree inferred from multiple alignment between known acidobacterial DsrD proteins. ML tree was inferred using PhyML v20160115 ran with following model and parameters: --pinv e --alpha e -- nclasses 4 -o tlr -f m --bootstrap 100. DsrD variants of Acidobacteria isolated from peats and soils are colored in green, of that from aquatic environment – in grey. *Archaeoglobus profundus* was used as the outgroup.

Dissimilatory sulfate reduction is known for Acidobacteriota members from metagenomic and metatranscriptomic studies. Acidobacterial species appear to be important drivers of C and S cycles in such hab tats as peats (Hausmann et al., 2018), thawing permafrost (Woodcroft et al., 2018), revegetated area near acid mine drainage (Li et al., 2022), and, as it was shown in this study, in the extremely contaminated Technosols. A recent study confirmed that Acidobacteria can couple degradation of complex plant biopolymers to sulfate reduction (Dyksma and Pester, 2023). Because of such metabolic preferences, these bacteria often encode multiple glycosyl hydrolases. In accordance, acidobacterial MAGs in this study also encoded different families of glycosyl hydrolases (supplementary material file S2, figure 4).

Several conditions could favor acidobacterial sulfate-reducers in Atamasnkoe Technosol. Firstly, acidobacteria can easily switch between anaerobic and aerobic metabolism, which is a useful trait in soils with regular flooding events. Secondly, due to the extreme level of contamination, hypoxic conditions, and overall metabolic suppression in Atamanskoe soils (Zamulina et al., 2021), slow-growing taxa (like Acidobacteriota) gain a selective advantage over other less-oligotrophic plant material-degraders. Finally, many acidobacterial classes are acidotolerant; correspondingly, 4 out of 6 acidobacterial MAGs in this study were assembled from low pH-soil metagenomes. Interestingly, marine derived DsrD variants in our analysis represented an inner clade relative to that of peat and soil habitats, suggesting the terrestrial origin of dissimilatory sulfur metabolism in Acidobacteriota.

### Atamanskoe sulfur oxidizers are remarkably multifunctional

A total of 31 MAGs (mostly, Pseudomonadota) encoded complete or almost complete SoxAB genes clusters (KEGG pathway reconstruction confirmed the full or one-block-missing Sox cluster). Among them, 14 MAGs also possesed dsrAB, and 9/14 – AprAB and sat proteins, but no DsrD or DsrD-like proteins. Moreover, ten DsrAB-containing bins also encoded DsrEFH, suggesting they can oxidize sulfur compound by both Sox and reverse-Dsr pathways (Neukirchen, Pereira and Sousa, 2023). Over half of SOB group also encoded a variety of organosulfonate monooxygenases and oxidoreductases. Moreover, the presence of sulfite oxidoreductases in their genomes (SUOX, SorA, SoeA, SoeB) suggests that they can utilize them to oxidize sulfite formed from organosulfonates. Several taxa showed incomplete sets of DMSO (dimethylsulfoxide) and DMSP (Dimethylpropiothetin) metabolism genes: bins 13_10, d4_12, 42_42, 42_6 and d3_36 could potentially transform DMSO from dimethylsulfone, while microbes under d2_14, 13_32_2, 13_37, d3_108, d3_99 and d2_76 bins can possibly transform metabolites of DMSP to Methylthiopropanoyl-CoA or methanethiol (which can in turn me metabolized by other member of community into sulfide or sulfite. Several recovered MAGs were affiliated with taxa rarely mentioned in the context of sulfur oxidation metabolism. *Pseudomonas yamanorum* (MAG 42_83) and the member of candidate Casimicrobiaceae family (MAG d2_57_2) were found to encode Sox genes, while Gemmatimonadales bacterium (MAG d3_36_2) genome, although possessed incomplete SoxAB cluster, contained SELENEBP1 and met genes. Alphaproteobacteria member (42_7) and *Pseudomonas yamanorum* contained 42 copies of tauD and 8 copies of ssuD correspondingly. Finally, MAGs d2_130_2 (Acidoferrales), d_019 (Limnocylindria), d4_19 (*Sulfuriferula multivorans*) and d2_36 (Terriglobia) can possibly oxidize thiosulfate via Dox pathway. Additional comments should be made for species under the d2_22, tr1_93 and d2_10 bins. Besides the complete set of Sox proteins, they encoded xsc and suyAB, producing sulfite out of sulfoacetaldehyde and sulfolactate, and also two independent enzymes capable of liberating sulfite moiety from methanesulfonic acid. Moreover, two of them (both betaproteobacteria) can possibly supply themselves with methanesulfonic using dimethylsulfone, since they have sfnG protein. This demonstrates profound multifunctionality and novelty of sulfur-metabolizing microbial community in Atamanskoe soils.

## Conclusions

The sulfur cycle exhibits immense diversity in terms of underlying taxa, metabolic reactions, and interconnections with other biogeochemical cycles. In recent years, researchers have begun to recognize that the terrestrial sulfur cycle differs from that of water environments. Our study aligns with this understanding and offers insights into the behavior of sulfur cycle bacteria in a unique soil environment of technogenic origin. We demonstrate that elevated concentrations of xenobiotic pollutants like HM and PAH, along with extreme levels of bulk sulfur content, low pH and fluctuating redox conditions, have given rise to a unique microbial community involved in sulfur transformation. This community is composed of: 1) known, but non-canonical taxa that carry out typical reactions, as seen with sulfate-reducing acidobacteria; 2) known taxa engaging in a multitude of oxidative, reductive, and organosulfur transformations, such as various sulfur oxidizing alpha-, gamma-, and betaproteobacteria; 3) previously unidentified taxa that have not been successfully cultured in a laboratory setting, like Binatota, Casimicrobiaceae, and Acidoferrales representatives, which may theoretically possess completely new genes associated with sulfur compound metabolism; 4) multiple taxa capable of collectively replenishing pools of readily consumable compounds, such as sulfite and thiosulfate. Consequently, these taxa contribute to the cryptic biogeochemical cycling of the compounds with high turnover rates but low absolute abundance. Furthermore, we demonstrate that sulfate-reducers – a significant part of microbial community according to culture-dependent analysis and the taxonomic profiling of metagenomes – can still represent only a small fraction of overall microbiome and require additional efforts to characterize it taxonomically and functionally. We conclude that sulfate-reducers indeed are members of soil rare biosphere and do not reach abundance comparable to that of aquatic habitats.

Studies on microbial sulfur cycles continue to uncover an inexhaustible source of new information, revealing the adaptive and diverse nature of microbial communities. It is important to acknowledge that we have only explored a small fraction of terrestrial ecosystems thus far. Therefore, further research in this field holds immense significance, both for practical and fundamental implications, such as mitigating greenhouse gas emissions and deepening our fundamental understanding of microbial behavior in nature.

