## Supplementary material file 2 for "Microbiota of the sulfur cycle in an extremely contaminated Technosol undergoing pedogenesis: A culture-dependent and metagenomic approach"

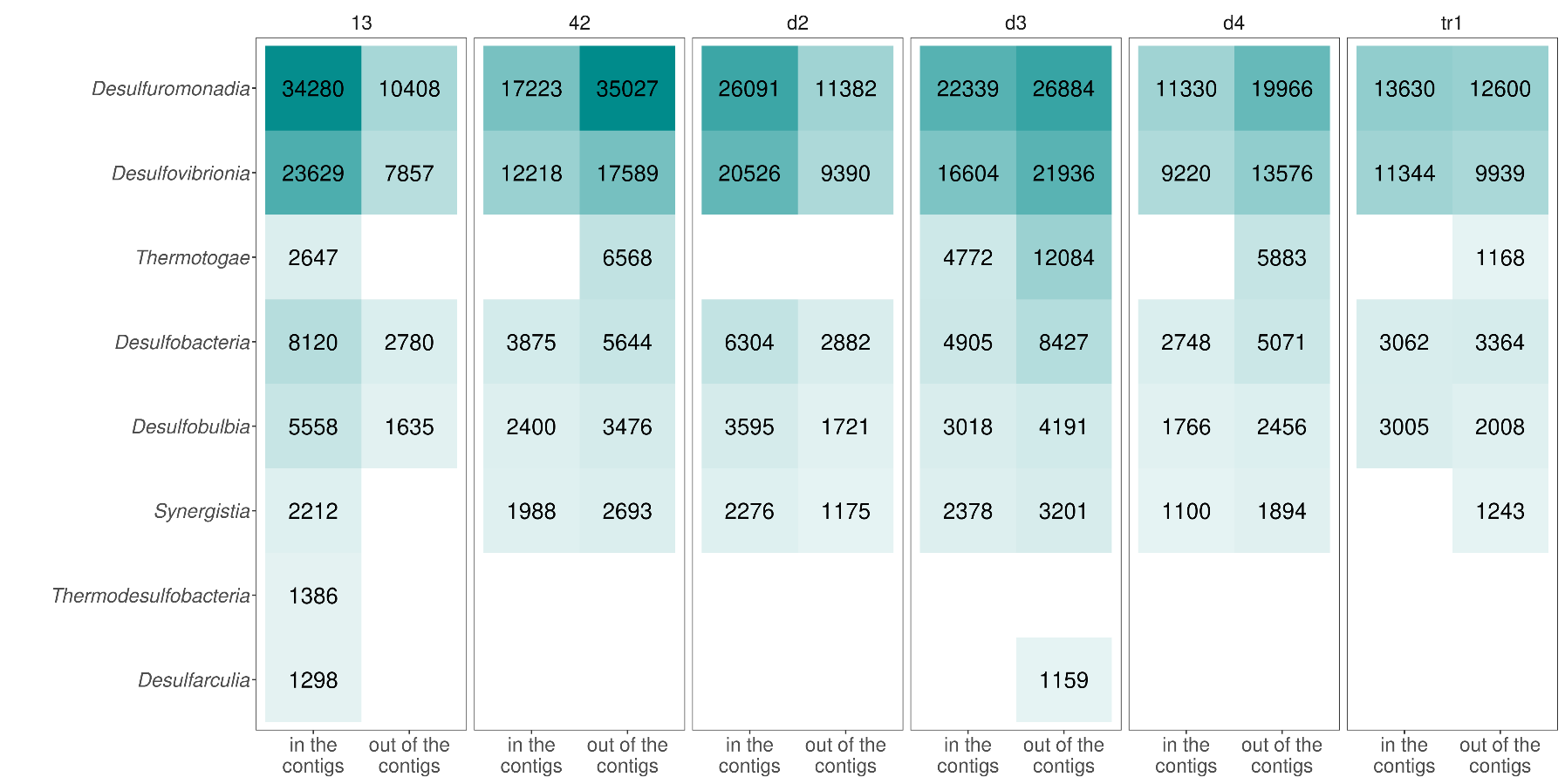

**Figure 1. Reads affiliated with different SRB classes that ended up either in contigs (and the subsequent bins) or out of them. Samples t6 and 28 were excluded based on the low SRB content in metagenomic profiles and culture-dependent data.**

**Acidobacterial DsrD tree coordinates in .nwk format:**

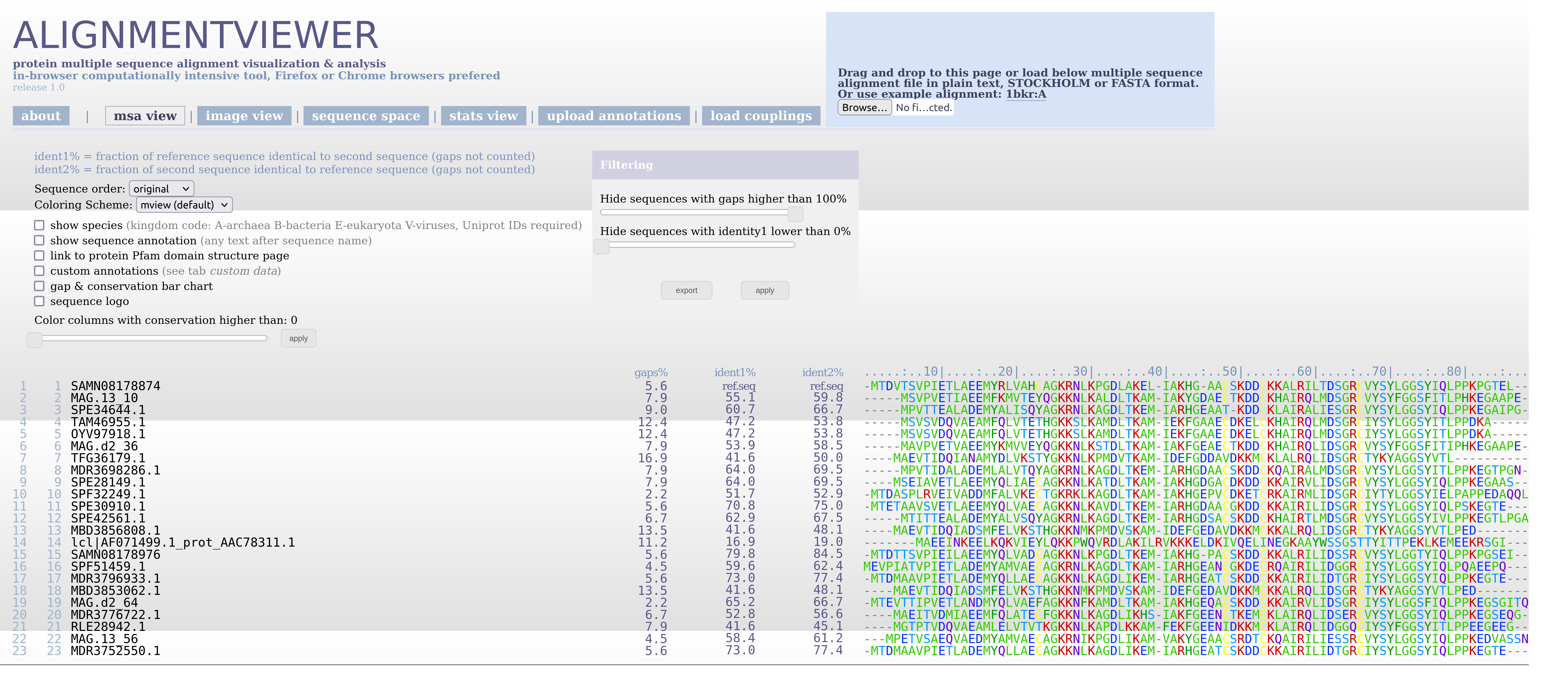
**Figure 2: Multiple alignment of DsrD proteins in all reported acidobacterial MAGs with Dsr genes.**

((((SPE30910.1:0.107252,((MDR3796933.1:1e-08,MDR3752550.1:1e-08)99:0.0753493,(SPE34644.1:0.096072,(MDR3698286.1:0.127411,SPE42561.1:0.0820724)38:0.0435672)85:0.276492)5:0.00451409)5:0.0227762,(SAMN08178976:0.0947589,SAMN08178874:0.103877)76:0.120008)13:0.0393626,((MAG.13_56:0.398729,(SPF32249.1:0.547302,SPF51459.1:0.248304)27:0.0508815)28:0.0875663,((SPE28149.1:0.147526,MAG.d2_64:0.213274)24:0.0168345,((MAG.13_10:0.0707061,MAG.d2_36:0.0760777)66:0.181672,(MDR3776722.1:0.289697,((TAM46955.1:1e-08,OYV97918.1:1e-08)93:0.169485,(RLE28942.1:0.300318,(TFG36179.1:0.0775468,(MBD3853062.1:1e-08,MBD3856808.1:1e 08)97:0.0714552)100:0.381338)93:0.0973969)67:0.163536)47:0.130141)34:0.175376)3:0.0408508)1:3.77e-06)100:3.000125365,lcl|AF071499.1_prot_AAC78311.1:4.066314635)1;

**DsrD sequences of known sulfate-reducing acidobacteria:**

>MAG.d2_36 [Terriglobia bacterium] This study

MAVPVETVAEEMYKMVVEYQGKKNLKSTDLTKAMIAKFGEAECTKDDCKHAIRQLIDSGRCVYSYFGGSFITIPHKEGAAPE

>MAG.13_10 [Terriglobia bacterium] This study

MSVPVETIAEEMFKMVTEYQGKKNLKALDLTKAMIAKYGDAECTKDDCKHAIRQLMDSGRCVYSYFGGSFITLPHKEGAAPE

>MAG.d2_64 [Terriglobia bacterium] This study

MTEVTTIPVETLANDMYQLVAEFAGKKNFKAMDLTKAMIAKHGEQACSKDDCKKAIRVLIDSGRCIYSYLGGSFIQLPPKEGSGITQ

>MAG.13_56 [Terriglobia bacterium] This study

MPETVSAEQVAEDMYAMVAECAGKRNIKPGDLIKAMVAKYGEAACSRDTCKQAIRILIESSRCVYSYLGGSYIQLPPKEDVASSN

>SPE30910.1 DsrD [Candidatus Sulfotelmatomonas gaucii] Peat soil

MTETAAVSVETLAEEMYQLVAECAGKKNLKAVDLTKEMIARHGDAACGKDDCKKAIRILIDSGRCIYSYLGGSYIQLPSKEGTE

>SPE28149.1 DsrD [Acidobacteriia bacterium SbA2]

MSEIAVETLAEEMYQLIAECAGKKNLKATDLTKAMIAKHGDGACDKDDCKKAIRVLIDSGRCVYSYLGGSYIQLPPKEGAAS

>SPE42561.1 DsrD [Candidatus Sulfopaludibacter sp. SbA3] Peat soil

MTITTEALADEMYALVSQYAGKRNLKAGDLTKEMIARHGDSACSKDDCKHAIRTLMDSGRCVYSYLGGSYIVLPPKEGTLPGA

>SPF51459.1 DsrD [Candidatus Sulfopaludibacter sp. SbA4] Peat soil

MEVPIATVPIETLADEMYAMVAECAGKRNLKAGDLTKAMIARHGEANCGKDECRQAIRILIDGGRCIYSYLGGSYIQLPQAEEPQ

>SPE34644.1 Dissimilatory sulfite reductase subunit D (DsrD) [Candidatus Sulfopaludibacter sp. SbA6] Peat soil

MPVTTEALADEMYALISQYAGKRNLKAGDLTKEMIARHGEAATKDDCKLAIRALIESGRCVYSYLGGSYIQLPPKEGAIPG

>SPF32249.1 DsrD [Candidatus Sulfotelmatobacter kueseliae] Peat soil

MTDASPLRVEIVADDMFALVKECTGKRKLKAGDLTKAMIAKHGEPVCDKETCRKAIRMLIDSGRCIYTYLGGSYIELPAPPEDAQQL

>OYV97918.1 SAMN06622430 MAG: hypothetical protein B7Z68_02315 [Acidobacteria bacterium 21-70-11]

MSVSVDQVAEAMFQLVTETHGKKSLKAMDLTKAMIEKFGAAECDKELCKHAIRQLMDSGRCIYSYLGGSYITLPPDKA

>TAM46955.1 SAMN10720137 MAG: hypothetical protein EPN53_12450 [Acidobacteria bacterium] Groundwater

MSVSVDQVAEAMFQLVTETHGKKSLKAMDLTKAMIEKFGAAECDKELCKHAIRQLMDSGRCIYSYLGGSYITLPPDKA

>RLE28942.1 SAMN09215175 MAG: hypothetical protein [Acidobacteria bacterium] Deep-sea hydrothermal vent sediments

MGTPTVDQVAEAMLELVTVTKGKKNLKAPDLKKAMFEKFGEENIDKKMCKLAIRQLIDGGQCIYSYFGGSYITLPPEEGEEG

>TFG36179.1 MAG: hypothetical protein [Candidatus Aminicenantes bacterium] Marine sediments

MAEVTIDQIANAMYDLVKSTYGKKNLKPMDVTKAMIDEFGDDAVDKKMCKLALRQLIDSGRCTYKYAGGSYVTL

>MBD3853062.1 MAG: hypothetical protein [Acidobacteria bacterium] Marine sediments

MAEVTIDQIADSMFELVKSTHGKKNMKPMDVSKAMIDEFGEDAVDKKMCKKALRQLIDSGRCTYKYAGGSYVTLPED

>MBD3856808.1 MAG: hypothetical protein [Acidobacteria bacterium] Marine sediments

MAEVTIDQIADSMFELVKSTHGKKNMKPMDVSKAMIDEFGEDAVDKKMCKKALRQLIDSGRCTYKYAGGSYVTLPED

>SAMN08178976 MAG. hypothetical protein [Ca. Acidiflorens stordalenmirensis] Thawing permafrost soil

MTDTTSVPIEILAEEMYQLVADCAGKKNLKPGDLTKEMIAKHGPACSKDDCKKALRILIDSSRCVYSYLGGTYIQLPPKPGSEI

>SAMN08178874 MAG. hypothetical protein [Acidiflorens clade 2] Thawing permafrost soil

MTDVTSVPIETLAEEMYRLVAHCAGKRNLKPGDLAKELIAKHGAACSKDDCKKALRILTDSGRCVYSYLGGSYIQLPPKPGTEL

>MDR3752550.1 MAG: hypothetical protein P4K78_01790 [Terracidiphilus sp.]

MTDMAAVPIETLADEMYQLLAECAGKKNLKAGDLIKEMIARHGEATCSKDDCKKAIRILIDTGRCIYSYLGGSYIQLPPKEGTE

>MDR3796933.1 MAG: hypothetical protein P4K93_02200 [Terracidiphilus sp.]

MTDMAAVPIETLADEMYQLLAECAGKKNLKAGDLIKEMIARHGEATCSKDDCKKAIRILIDTGRCIYSYLGGSYIQLPPKEGTE

>MDR3776722.1 MAG: hypothetical protein P4K97_07485 [Terracidiphilus sp.]

MAEITVDMIAEEMFQLATECFGKKNLKAGDLIKHSIAKFGEENCTKEMCKLAIRQLIDSERCVYSYLGGSYIQLPPKEGSEQG

>MDR3698286.1 MAG: hypothetical protein P4L56_01545 [Candidatus Sulfopaludibacter sp.] Peat from acidic fen

MPVTIDALADEMLALVTQYAGKRNLKAGDLTKEMIARHGDAACSKDDCKQAIRALMDSGRCVYSYLGGSYITLPPKEGTPGN

>lcl|AF071499.1_prot_AAC78311.1_4 [gene=dsrD] [Archaeoglobus profundus]

MAEEINKEELKQKVIEYLQKKPWQVRDLAKILRVKKKELDKIVQELINEGKAAYWSSGSTTYITTPEKLKEMEEKRSGI

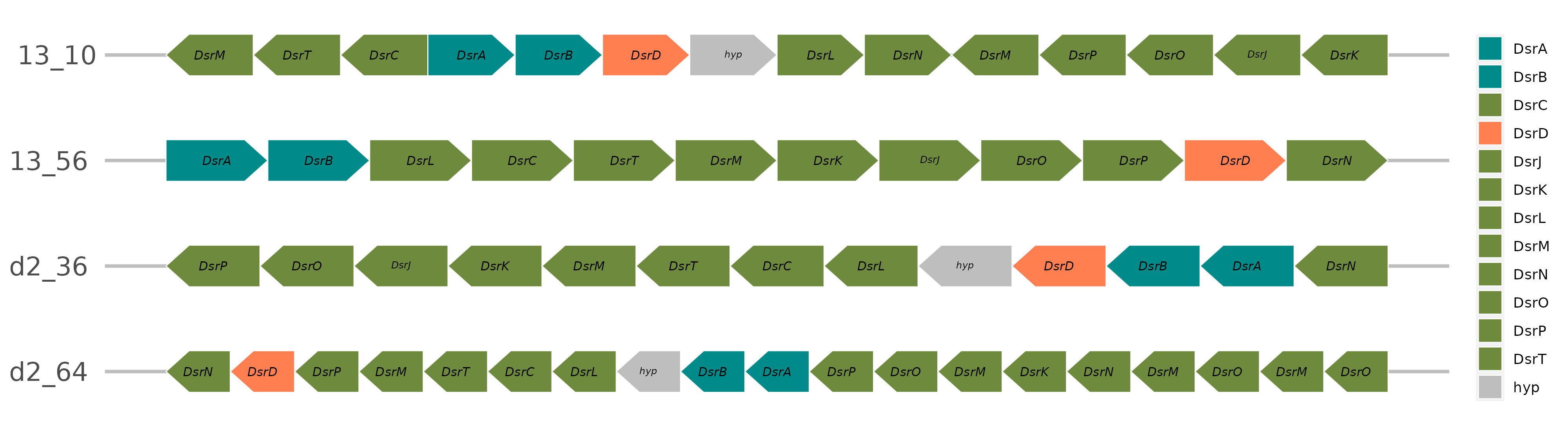
**Figure 3: Schematic representation of Dsr operon structures in four Terriglobia bins assembled from Atamanskoe technosol metagenomes. "hyp" refers to DsrAB-associated protein with unknown function (based on BLASTp inference).**

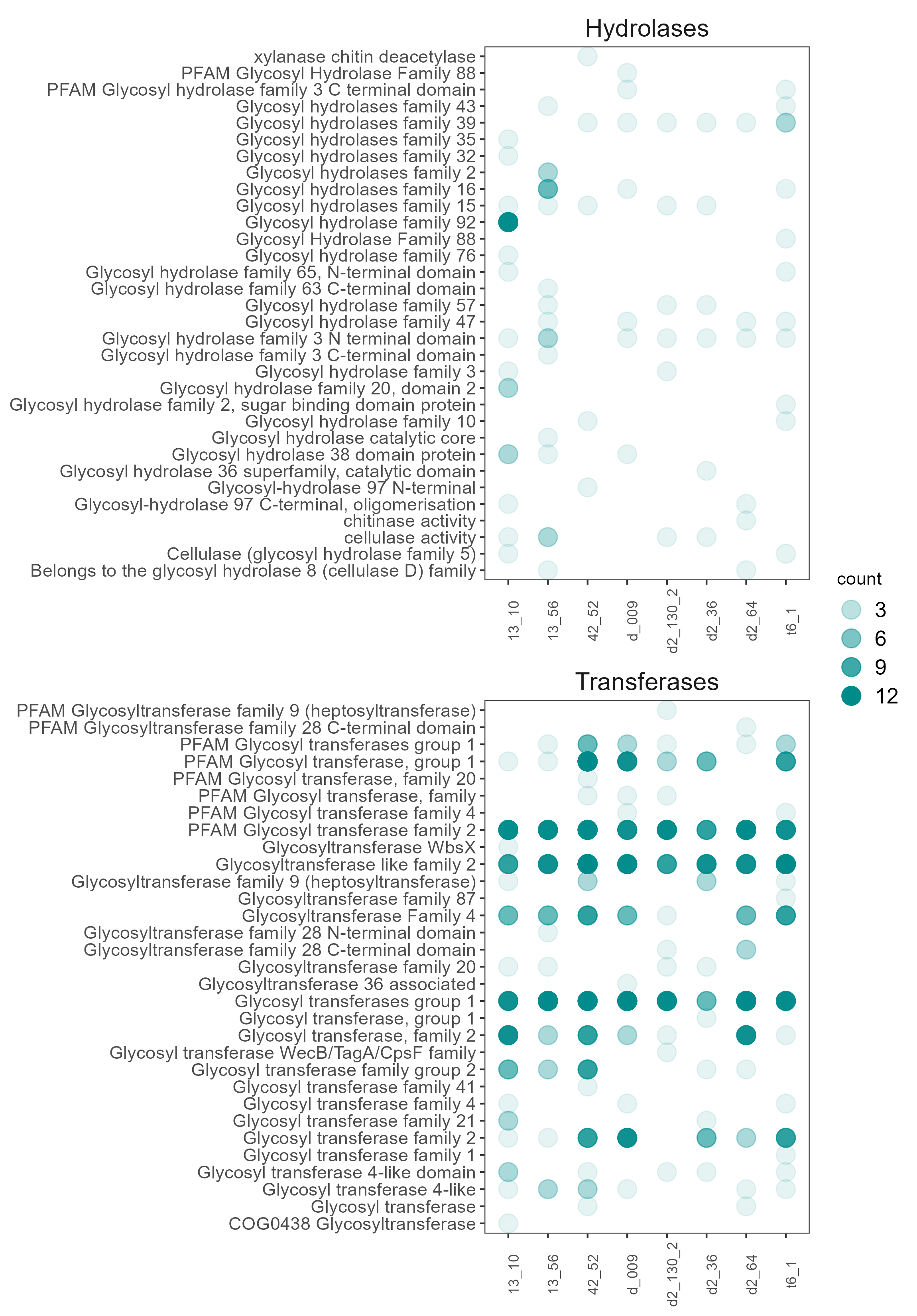

**Figure 4: Glycosyl Hydrolases and transferases families identified in eight acidobacterial bins.**
